# Linking cognitive integrity to working memory dynamics in the aging human brain

**DOI:** 10.1101/2023.08.18.553840

**Authors:** G Monov, H Stein, L Klock, J Gallinat, S Kühn, T Lincoln, K Krkovic, PR Murphy, TH Donner

**Author notes:** These authors contributed equally to this work (PR Murphy, TH Donner). Correspondence to: Gina Monov, Full address: Section of Computational Cognitive Neuroscience, Department of Neurophysiology & Pathophysiology, University Medical Center Hamburg-Eppendorf, Martinistraße 52, 20246 Hamburg, Germany.

## Abstract

Aging is accompanied by a decline of multiple cognitive capacities, including working memory: the ability to maintain information online for the flexible control of behavior. Working memory involves stimulus-selective neural activity, persisting after stimulus presentation in widely distributed cortical areas. Here, we unraveled the mechanisms of working memory in healthy older adults and patients with mild cognitive impairment (MCI), a condition associated with increased risk of developing dementia. We studied a sample of 19 older adults diagnosed with MCI and 20 older healthy controls using a combination of model-based behavioral psychophysics, neuropsychological assessment, and magnetoencephalographic (MEG) recordings of brain activity. Twenty-one younger healthy adults were studied with model-based behavioral psychophysics only. All subjects performed a visuo-spatial delayed-match-to-sample working memory task under systematic manipulation of the temporal delay and the spatial distance between successively presented sample and test stimuli. We developed a computational model of the latent dynamics underlying task behavior and fit this to individual behavior. In the model, working memory representations diffused over time, a threshold was applied to produce a match/non-match decision about sample and test locations, and occasional lapses produced random decisions. For the older participants, we related the individual model parameters to a summary measure of individual cognitive integrity obtained from a large neuropsychological test battery, as well as to cortical MEG activity during the delay interval of the working memory task. For all groups, task accuracy decreased with delay duration and sample-test distance. When sample/test distances were small, older adults exhibited larger false alarm rates than younger adults. The behavioral effects were well captured by the model, which explained the age-related differences in terms of a deterioration of the quality of working memory representations, rather than differences in task strategy (i.e., threshold parameter). Task accuracy as well as the parameters governing behavioral stochasticity (diffusion noise and lapse rate combined) were correlated with overall cognitive integrity in the MCI group, but not in the older healthy controls. Individual task accuracy and stochasticity parameters (diffusion noise and lapse rate) were also correlated with stimulus-selective cortical activity during the delay interval, as assessed by decoding of the MEG signals, corroborating their validity as markers of cortical working memory mechanisms. Our findings provide insight into the mechanistic basis of aging-related changes in working memory maintenance and reveal a link between individual working memory dynamics and cognitive integrity in MCI.

## Introduction

Aging has profound effects on higher brain function, with important ramifications at the levels of society and individuals.^1^ Several cognitive capacities tend to decline during healthy aging in a largely correlated fashion.^1,2^ At the same time, cognitive aging exhibits strong differences between individuals.^1^ These individual differences may reflect undetected pathology of neural circuitry as well as idiosyncratic strategies to cope with performance reductions in specific tasks.^2^ Developing a mechanistic understanding of these age-related changes in cognition faces several challenges.^2^ One challenge is to delineate factors that exacerbate the physiological age-related cognitive decline and incur an increased risk of developing dementia, as manifested for example in MCI.^2–4^ Another challenge is to isolate specific mechanistic markers of aging-related changes in cognitive computation, distinct from compensatory strategies, that predict individuals’ performance on specific cognitive tasks as well as their general cognitive integrity.^2^ Our current study aims at mastering these challenges.

One important capacity affected by age-related cognitive decline is working memory.^1–3,5^ Working memory refers to the ability to maintain information online and put this information to use for cognitive computation and action.^6^ Working memory is an appealing focus for unraveling the physiological basis of age-related cognitive decline for several reasons. First, working memory is a fundamental building block of cognition^7,8^, and so individual working memory performance tends to predict performance on a variety of other cognitive tasks^e.g. 9–12^. Second, the neural basis of the active maintenance of information, including even age-related deteriorations^13^, has been studied extensively in the monkey brain as well as in cortical circuit models^8,14^. Third, recent advances in neuroimaging data analysis now also permit non-invasive tracking of the active maintenance of working memory content in the human brain.^15,16^

Previous work on working memory has culminated in a cohesive framework for linking performance on working memory tasks to computational and neurophysiological mechanisms. The maintenance of information in working memory relies, at least in part, on stimulus-selective neuronal activity that persists after the offset of a to-be-remembered stimulus.^7,8,14^ Convergent evidence from different approaches in humans and monkey has identified such content-selective persistent activity in many brain regions including frontal and parietal association cortex as well as even sensory cortex.^8,14,15,16,17–19^ Modeling work shows that such persistent activity can be produced by synaptic reverberation^14,20,21^ in circuits with a balanced interplay between recurrent excitation and inhibition^22^. Disruptions in this balance deteriorate the stability of the stimulus-selective activity patterns, thus yielding a characteristic decrease of behavioral performance with the duration of working memory delays.^23^

This framework now opens the door for studying the neural bases of age-related changes in cognition at an unprecedented level of mechanistic detail. Importantly, abstractions of the above (high-dimensional and biophysically detailed) cortical circuit models can be fit to behavioral data and used to decompose an individual’s performance in a working memory task into several latent sources of behavioral variability.^24–27^ This enables a dissociation of the stability of the working memory representation per se from task strategies, thus helping to pinpoint the sources of variability in task performance.

We set out to identify dynamical mechanisms underlying age-related changes in cognition. We aimed to (i) identify computational and neurophysiological signatures of working memory mechanisms and (ii) link these changes to individual levels of cognitive integrity in older adults. To these ends, we combined model-based analyses of behavior during a working memory task with MEG recordings of cortical population activity. We isolated specific signatures of the stability (i.e., quality) of working memory representations, and the underlying persistent cortical activity during working memory delays, from strategic factors that also affect working memory performance and are malleable by aging.

## Materials & methods

### Participants and recruitment

We report analyses of two datasets acquired in the context of different studies. Participants of both studies were financially compensated with ten Euros per hour of participation and (for dataset 2) for expenses incurred due to SARS-CoV-2-antigen testing.

### Sample for dataset 1

Dataset 1 included behavioral (working memory task), neuropsychological, and MEG data from three groups of older participants (Fig. 1A and Table 1): Participants diagnosed clinically with Mild Cognitive Impairment (MCI; N=19), age-matched older healthy controls (OHC; N=20), and unclassified older participants (UNC; N=7). The recruitment of participants and definition of these groups is described in the following.

**Figure 1.**
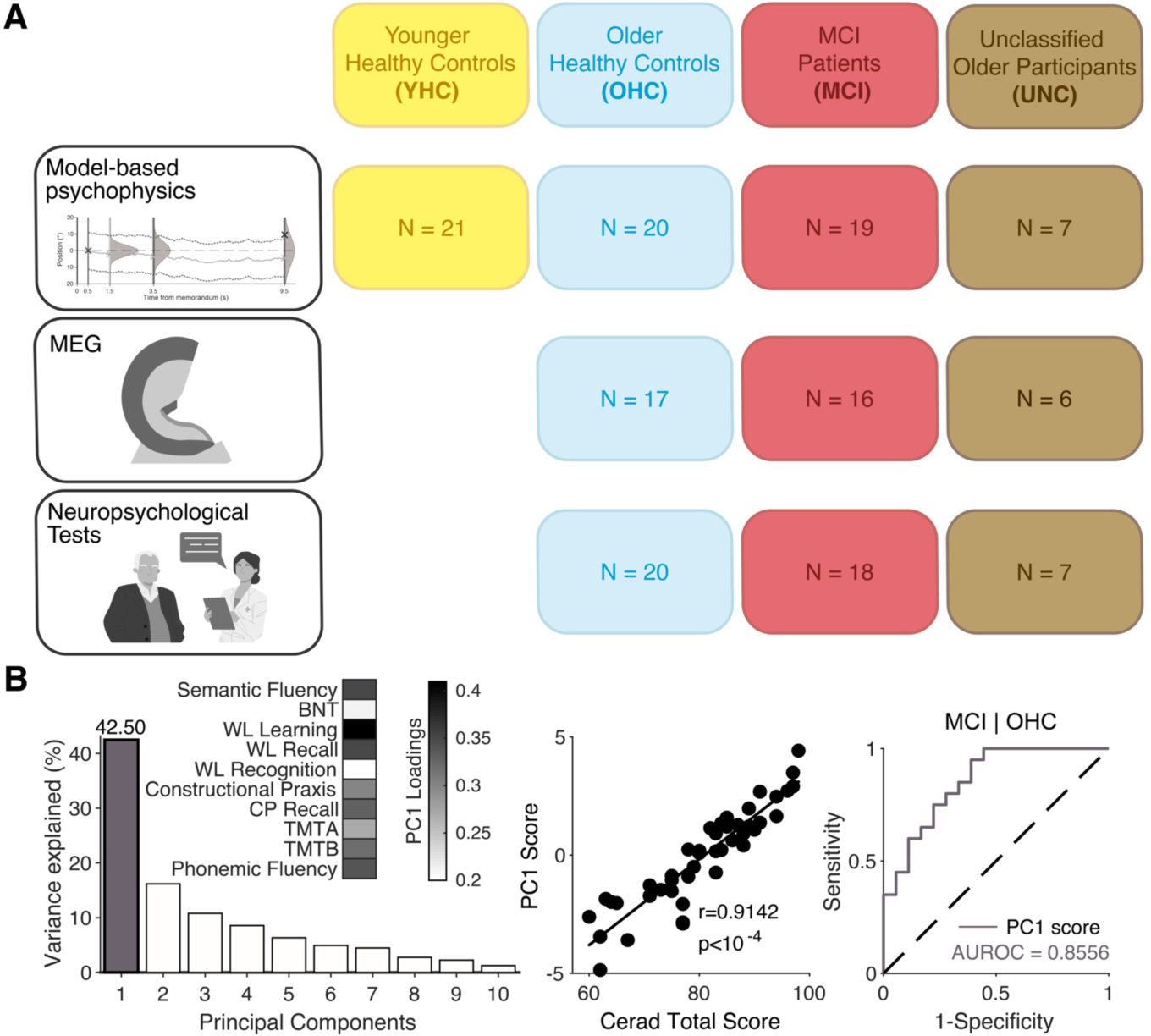
Sample and approach. **(A)** Testing modalities (rows) and definition of participant subgroups (columns). Sample sizes within each modality/subgroup combination defines the number of subjects included in the corresponding analyses. **(B)** *Left*: Variance explained by the principal components derived from PCA of the CERAD-Plus neuropsychological test battery data. Inset shows loadings of different tests for the first principal component (PC1, grey in plot of variance explained). *Middle*: Correlation of PC1 scores and CERAD total scores across all older adults. *Right*: Receiver operating characteristic (ROC) curves for diagnostic classification (OHC vs. MCI) using PC1 scores (grey).

**Table 1.** Sample description.

| *Measure* | **YHC (N=21)** | **OHC (N=20)** | **MCI (N=19)** | **UNC (N=7)** |
| --- | --- | --- | --- | --- |
| *Mean age in years (S.D.)* | 27.2 (3.8) | 69.4 (6.2) | 73.7 (6.2) | 70.3 (8.2) |
| *% Female* | 66.7 | 60 | 63.2 | 57.1 |
| *% Diagnosis: amnesic MCI \| multidomain-amnesic MCI* | - | - | 63.2 \| 36.8 | - |

Participants for the MCI group were recruited in the outpatient clinic for memory disorders of the Department for Psychiatry and Psychotherapy, University Medical Center Hamburg-Eppendorf (UKE). The study was part of a longitudinal study, in which patients underwent a placebo-controlled intervention (tailored video game). All measurements reported in this paper stem from the first time point (pre-intervention baseline). Included patients fulfilled diagnostic criteria for MCI (F06.7) according to the International Classification of Diseases, Version 10^28^. Further, the domain-specific subdivisions of amnestic MCI and multidomain-amnestic MCI were both eligible for the study. The distribution of diagnoses within the MCI group is shown in Table 1.

Age-matched healthy control participants for the OHC group were recruited using flyers and newspaper/online announcements. For the assessment of cognitive integrity, the extended version of the well-validated Consortium to Establish a Registry for Alzheimer’s Disease (CERAD-Plus) neuropsychological test battery including additional tests for executive functions and cognitive speed^29–31^ was employed. Participants of the OHC group showed no cognitive impairment unexpected for their age, gender, and educational level (population-based z-scores greater than −1.5 in each test of the test battery).

Participants originally recruited for the OHC group that produced scores in line with MCI in the neuropsychological assessment could choose to be consulted clinically by a psychiatrist of the outpatient clinic for memory disorders. In case they met the inclusion criteria for the study and the physician deemed it appropriate, participation as an MCI patient in the study was offered and they were assigned to the MCI group for further analysis (N=1). Those subjects who did not fulfill the criteria for OHC and did not undergo this full diagnostic procedure were not assigned to either the OHC or MCI group and were grouped together as the unclear diagnosis (UNC) group (N=7).

General inclusion criteria were participant’s age between 55 and 90 years, ability to consent, place of residence around Hamburg, Germany, and sufficient mobility. Exclusion criteria were: relevant psychiatric concomitant diseases (depression, schizophrenia, anxiety disorder, personality disorder), physical illnesses with a relevant influence on mental or motor skills functions (e.g. stroke, heart failure, cerebrovascular diseases, endocrinological disorders, inflammatory diseases of the central nervous system, epilepsy, Parkinson’s disease), substance dependence or substance abuse, relevant impairments of the sensory system that make interventions impossible, clinically relevant anemia, non-removable metal implants or implanted electronic devices and claustrophobia.

### Sample for dataset 2

Dataset 2 consisted of behavioral data (same working memory task as dataset 1) from a group of young healthy controls (YHC, N=21; Fig. 1A and Table 1), collected in the context of a separate study at the Department of Clinical Psychology and Psychotherapy at the Universität Hamburg. We used these data as a reference in the current study to identify overall age-dependent changes in working memory performance.

Participants in the YHC group were recruited from the general population of Hamburg through flyers and online announcements and included if they were between 18 and 35 years old and had no pre-existing, diagnosed mental disorder or neurological condition.

### Informed consent

Participants’ informed (written) consent was obtained in both studies. The study design for both datasets was approved by the ethics committee of the Department of Psychiatry at the University Medical Center Hamburg-Eppendorf (dataset 1) or the Faculty of Psychology and Human Movement Science at Universität Hamburg (dataset 2) and conducted in accordance with the Declaration of Helsinki. We excluded subjects from all analyses if their accuracy in the working memory task was below 60% (N=3; Supplementary Fig. 1C), which is below the accuracy that would be achieved should a participant employ the simple strategy of giving a “different” response on every trial; see ‘Working memory task and procedure’).

## Data acquisition and experimental setup

### Dataset 1

MEG sessions were conducted in the UKE Department of Neurophysiology and Pathophysiology. For MEG data acquisition, subjects were placed in a comfortable position in a magnetically shielded room, at a viewing distance of 51 cm from the screen on which the stimuli were shown. MEG data were recorded with 275 axial gradiometers (CTF Systems) at a sampling rate of 1200 Hz. ECG, vertical EOG, and horizontal EOG were measured using bipolar Ag/AgCl electrodes and a ground measured on the wrist. Subjects were asked to minimize their head movements during the measurements, and three-dimensional head position was recorded via fiducial coils attached to the external auditory canals and the nasion. This permitted online tracking of the head position and guiding of subjects back into their initial position during breaks of the experimental task. Stimuli were backprojected on a transparent screen with a projector (Sanyo PCL-XP51) with 1920 × 1080 resolution, at a refresh rate of 60 Hz. Eye movements and pupil size were recorded during task performance with an EyeLink® 1000 Long Range Mount (SR Research) at a sampling rate of 1000 Hz.

Structural T1-weighted MRI scans for individualized source reconstruction of MEG data were collected with a Siemens 3-Tesla MAGNETOM Prisma scanner (Erlangen, Germany) using a standard 32-channel head coil. The structural images were obtained using a three-dimensional T1-weighted magnetization prepared gradient-echo sequence (MPRAGE) (repetition time = 2500 ms; echo time = 2.12ms; TI = 1100 ms, acquisition matrix = 232 × 288 × 19.3, flip angle = 9°; 0.83 × 0.83 × 0.94 mm voxel size).

The neuropsychological assessment of the older subjects was administered by experienced clinical staff in the outpatient center for memory disorders of the UKE Department of Psychiatry and Psychotherapy. The tests of the CERAD-Plus battery were performed in a paper-pencil format and evaluated for diagnostic purposes according to the reference data provided by the Memory Clinic of Basel^32^.

### Dataset 2

Data were collected in a behavioral laboratory under similar conditions as for the MEG recordings in dataset 1 (task-relevant stimulus parameters were identical with the exception of a shorter inter-trial interval, see ‘Working memory task and procedure’). Here, we used a headrest to ensure a fixed viewing distance of 52 cm from the monitor, and stimuli were presented on a 22” Dell P2210 monitor with a resolution of 1680 × 1050 and refresh rate of 60 Hz. Eye-tracking (SMI RED500) and electrocardiogram (ECG) data were acquired during task performance but are not reported here.

### Neuropsychological test battery and cognitive integrity score

The neuropsychological diagnostics carried out were based on the validated CERAD-Plus test battery, which consists of eleven individual tests: verbal fluency (semantic: animals and phonemic: S-words), modified Boston Naming Test (BNT), Mini-Mental State Examination (MMSE), word list (learning, recall and recognition), constructional praxis, constructional praxis recall, and parts A and B of the Trail Making Test (TMT). As is common in the field, performance on the word list recognition test was quantified in terms of the discriminability score introduced in Mohs et al.^33^:

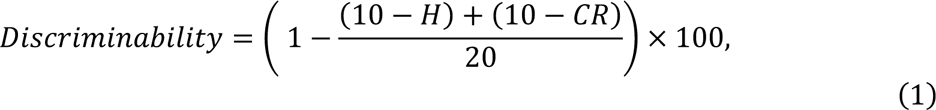

where H was the hit rate and CR was the rate of correct rejects.

We used principal component analysis (PCA) to combine ten test scores into a single summary measure of cognitive integrity, whereby the MMSE was excluded as scores on this test already reflect a mixture of assessments of several cognitive domains. The strong correlation between MMSE scores and our cognitive summary score (*r* = 0.65, *p*<10^−4^) further validated our approach (Supplementary Fig. 1B). To compute the summary measure of cognitive integrity, the individual scores on each test were z-transformed across subjects (Supplementary Fig. 1A). Unlike the other tests, high scores of both Trail Making Tests (parts A and B) reflect poor performance. To ensure that positive values reflected higher performance for all tests, the signs of the z-scores for these two tests were flipped. We then computed the (across-subjects) covariance matrix between the z-scored test scores (dimensionality: 10 × 10) and used MATLAB’s (MathWorks®) singular value decomposition algorithm to compute the corresponding ten principal components and their associated eigenvalues. One subject (MCI) was excluded from this analysis because they did not complete all tests in the battery.

For comparison to previous neuropsychological work, we also computed a composite CERAD total score proposed by Chandler et al.^34^ as the sum of the following six test scores: semantic fluency (max. 24 points), Boston Naming Test, word list learning, word list recall, word list recognition (*true positives* − *false positives*) and constructional praxis. This CERAD total score was closely correlated with the eigenvalue of the first principal component derived via the PCA procedure described above (Fig. 1B, *middle*). In this study, we used the latter as the summary measure of individual cognitive integrity, because it used information from all tests which proved to be diagnostically beneficial^31^ and explained a large fraction of variance (>40%) in the neuropsychological data (see ‘Results’).

### Working memory task and procedure

The visuospatial delayed match-to-sample working memory task (Fig. 2A; Supplementary Fig. 2A) was programmed in MATLAB using Psychtoolbox-3 ^35^. The task was to decide whether a sample stimulus and a test stimulus separated by a variable delay occurred in the same or different locations. Each trial began with presentation of a central white fixation cross (arm length: 0.8 degrees of visual angle, d.v.a.; arm thickness: 0.2 d.v.a.) that was present for the entire trial. After a variable baseline interval (uniform distribution with range 1-2.5s for dataset 1, 0.5-2.0s for dataset 2), the sample stimulus was presented for 0.5s, followed by the delay (1, 3 or 9s, equiprobable) and then the test stimulus (0.5s). Sample and test stimuli were circular checker-board patches (diameter: 2.8 d.v.a.; spatial frequency: 1 cycle per d.v.a), appearing in the lower visual hemifield at a fixed eccentricity of 6 d.v.a.. The sample could be presented at any of 12 equiprobable locations, ranging from ∼13.85° to ∼166.15° of polar angle (fixed spacing ≈ 13.85°), while the most extreme samples could still be flanked by a ‘near non-match’ test stimulus on both sides (amounting to 14 possible test stimulus locations, ranging from 0° to 180°). Herein, 0° refers to the left part of the horizontal meridian. The test occurred at either the same location as the sample or at a different location (see below). Upon offset of the test stimulus, the fixation cross changed color from white to light blue, which prompted subjects to report their decision via right-or left-handed button press for “same” or “different” judgments, respectively. This response was soon (0.1s) followed by visual feedback about its accuracy (“Correct” in green font; “Error” in red font; font size 36, presented 1.0 d.v.a. above fixation for 0.75s). Each trial was followed by a fixed interval (3s for dataset 1, 2s for dataset 2) during which participants were instructed to blink if needed, and this was followed by the baseline period of the following trial.

**Figure 2.**
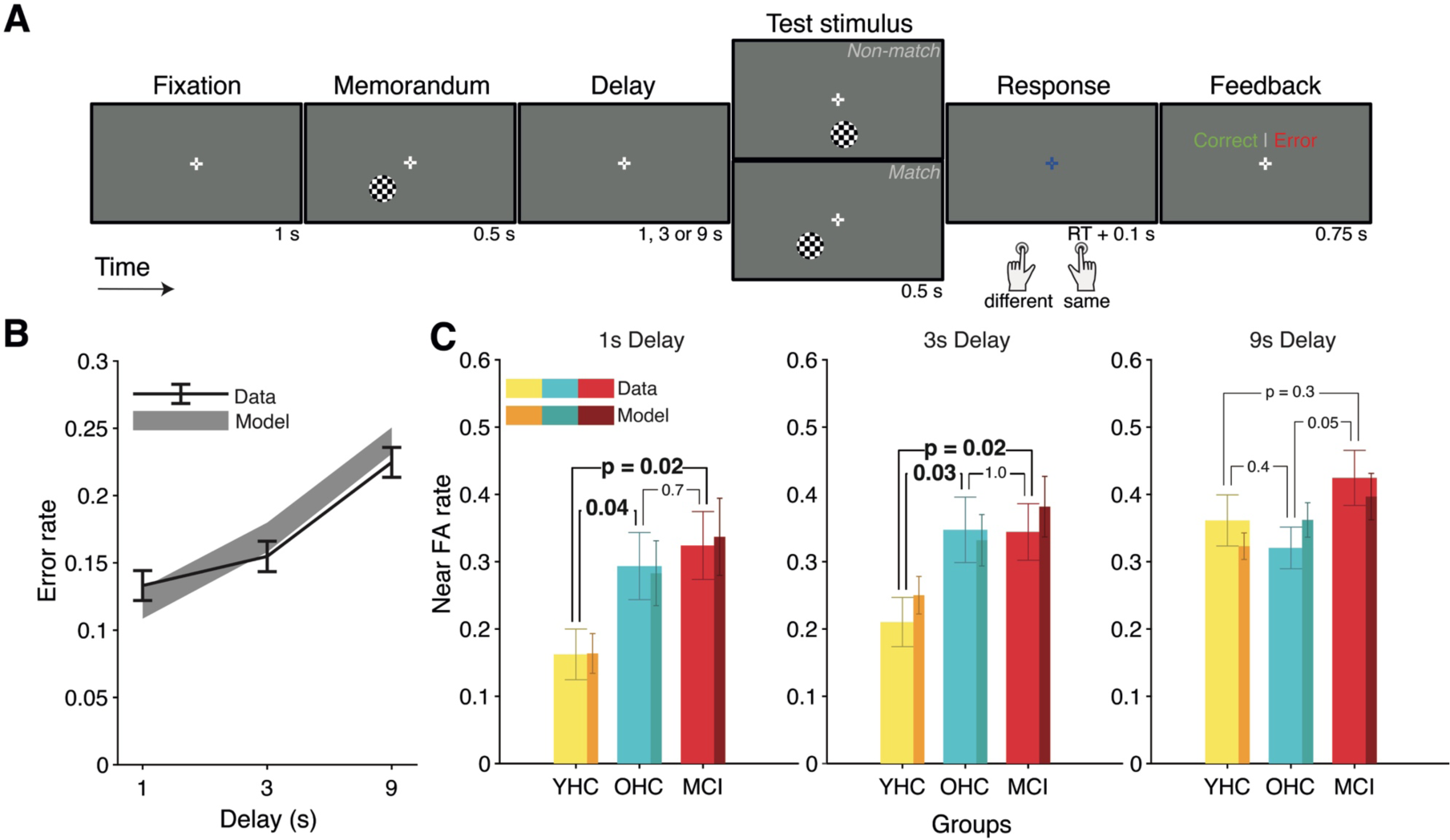
Working memory task and behavioral results. **(A)** Schematic of the delayed-match-to-sample working memory task. Stimuli shown are for illustration purposes only; for original appearance, see Supplementary Fig. 2A. **(B)** Error rate (black, mean ± s.e.m.) as a function of delay duration across all subjects (N=67) and corresponding model predictions (grey, mean ± s.e.m.). **(C)** False alarm rate on near non-match trials by delay durations (lighter shades, mean ± s.e.m.) and corresponding model predictions (darker shades, mean ± s.e.m.) for YHC (N=21; yellow), OHC (N=20; blue) and MCI (N=19; red) subgroups. P-values refer to results of non-parametric, between-subjects permutation tests.

The task was designed to consist of three trial categories, each with a desired frequency of occurrence within a block of 63 trials: ‘Match trials’ (sample and test at identical positions, 33% of trials), ‘Near non-match trials’ (smallest possible sample-test distance of 13.85°; 33% of trials), and ‘Far non-match trials’ (sample-test distance randomly chosen from the remaining possible sample-test distances, which could be between 27.7° and ∼166.15° depending on the sample location; 34% of trials). Trials were presented in blocks of 63 trials each, within which the different delay durations and sample-test distances were randomly interleaved under the above-mentioned constraint. Subjects received feedback about their average performance at the end of each block. They were instructed to fixate the central cross and minimize blinking during the trial.

Before starting the experimental task, all subjects underwent training to familiarize them with the task. This consisted of a general instruction of the task rules with the help of a slide presentation (administered outside the MEG chamber for dataset 1 and in the testing room for dataset 2) and practice with various aspects of the task (after the subject had been placed in the MEG for dataset 1, and in the testing room for dataset 2). The first stage of the practice required the participant to fixate the fixation cross while checkerboard stimuli identical to those of the main experiment were presented, including feedback if and when the participant broke fixation. Next, four non-consecutive example trials covering match, far non-match and near non-match trials and varying delay durations were performed. In case of an incorrect response on any of these trials, a text with the correct solution was displayed and the trial was repeated. Finally, six consecutive trials were performed with identical timing and inter-trial intervals as in the main experiment. Once training was complete, each subject then performed several blocks of the main experiment (concurrent to MEG measurement for dataset 1).

We aimed for three task blocks per subject in dataset 1 and at least two blocks per subject in dataset 2. Data collection had to be terminated early in some subjects, due to lack of alertness or willingness to continue, or end of the scheduled testing session. We obtained the complete set of three trial blocks for the following fractions of participants per group: MCI: 14/19; OHC: 13/20; UNC: 3/7; YHC: 4/21. Since the differences in trial counts for different individuals/groups studied here only affect the precision of the parameter estimates (behavioral model parameters or MEG measures), but did not bias them in a particular direction, they also did not bias the group comparisons or across-subjects correlations reported in this paper. Furthermore, we included these trial counts as nuisance regressors in all regression models.

### Analysis of working memory-guided behavior

Trials were excluded from analysis contingent on the following criteria: a task-irrelevant button (2 of 4 available buttons on the response pad) was pressed, response time was ≤ 0.2 s, or it exceeded the subject’s mean response time by 4 standard deviations. Further, the entire first block of one MCI participant was excluded from analysis due to poor comprehension of task instructions (accuracy <50 % correct for this block). Accordingly, an average of 157 trials per subject were submitted to analysis (range across subjects: 59 to 189 trials).

Mean response accuracy was computed as the proportion of correct responses (i.e., “same” response on match trials and “different” response on non-match trials). For several analyses (Fig. 2C, Supplementary Fig. 2A), error rates were analyzed separately for the three trial categories described above.

We also quantified the signal-detection theoretic^36^ measures of sensitivity (*d*′) and criterion (*c*) from the fractions of hits (“same” responses on match trials, denoted as *H*) and false alarms (“same” responses on non-match trials, denoted as *FA*), whereby the latter fraction was first computed separately for the near and far non-match trials and then averaged.

Sensitivity *d*′ and criterion *c* were then computed as follows:

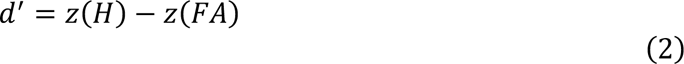

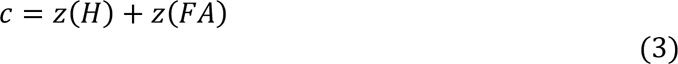

## Computational modeling of working memory-guided behavior

### General approach

Our model of working memory task behavior consisted of two computational elements, both of which accounted for a fraction of behavioral imprecision: (i) a point-estimate memory representation that diffused over time, leading to an increase in error as a function of delay^e.g. 25^; and (ii) a decision transformation of that representation into a categorical behavioral report ^e.g. 23,37^.

The dynamics of the memory representation were modelled as a Wiener diffusion process where the standard deviation of the across-trial distribution of memory representations at time *t* during the memory delay was captured by σ*_t_* and determined by the memory noise parameter

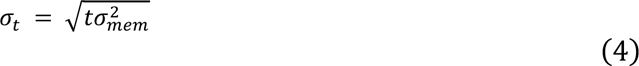

The across-trial variance of this memory representation thus increased monotonically as a function of time (eq. 4). This representation was transformed into a probability distribution over the task-relevant decision variable *x*, the absolute distance between the memory representation and test stimulus presented at the end of the delay of duration *T*. We modeled the decision variable as a normal distribution with a mean equal to the true sample-test distance (Δ), and folded it around zero (corresponding to test location) to reflect the absolute deviation of the internal memory representation from the test stimulus, as follows:

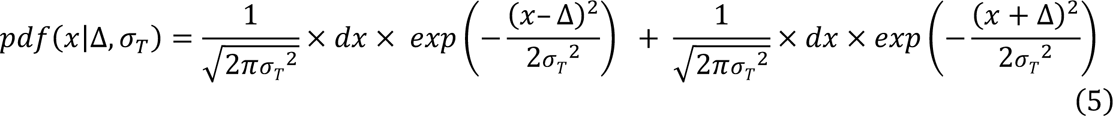

We modelled the decision function (*DF*) for a given value of *x* as a logistic function:

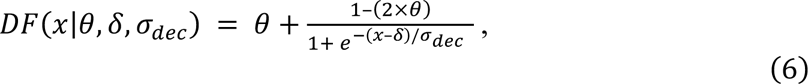

where δ was the inflection point of the function (*DF* (δ) = 0.5) and corresponds to a ‘soft’ (i.e. non-deterministic) threshold for translating *x* into a same (*DF* << 0.5) or different (*DF* >> 0.5) choice; σ*_dec_* was a decision noise parameter that governed the slope of *DF*; and θ was the probability of a time-independent lapse that determined the function’s two asymptotes (θ and 1-θ, respectively), assumed to be symmetric for simplicity. The probability of a “different” response as a function of delay duration *T* and sample-test distance Δ was then computed numerically, by integrating (i.e., summing) over all *x* for 0 ≤ *x* ≤ 360, and *dx* = 0.05:

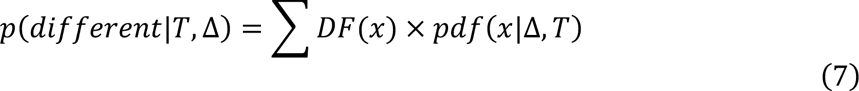

Note that lapses could be sensory, motor, decisional or mnemonic in origin. Also note that, if DF took the form of a step-function (i.e., infinite slope, σ*_dec_* = 0), then the location of the step was equivalent to a deterministic threshold applied to the decision variable (Fig. 3A, *middle*).

**Figure 3.**
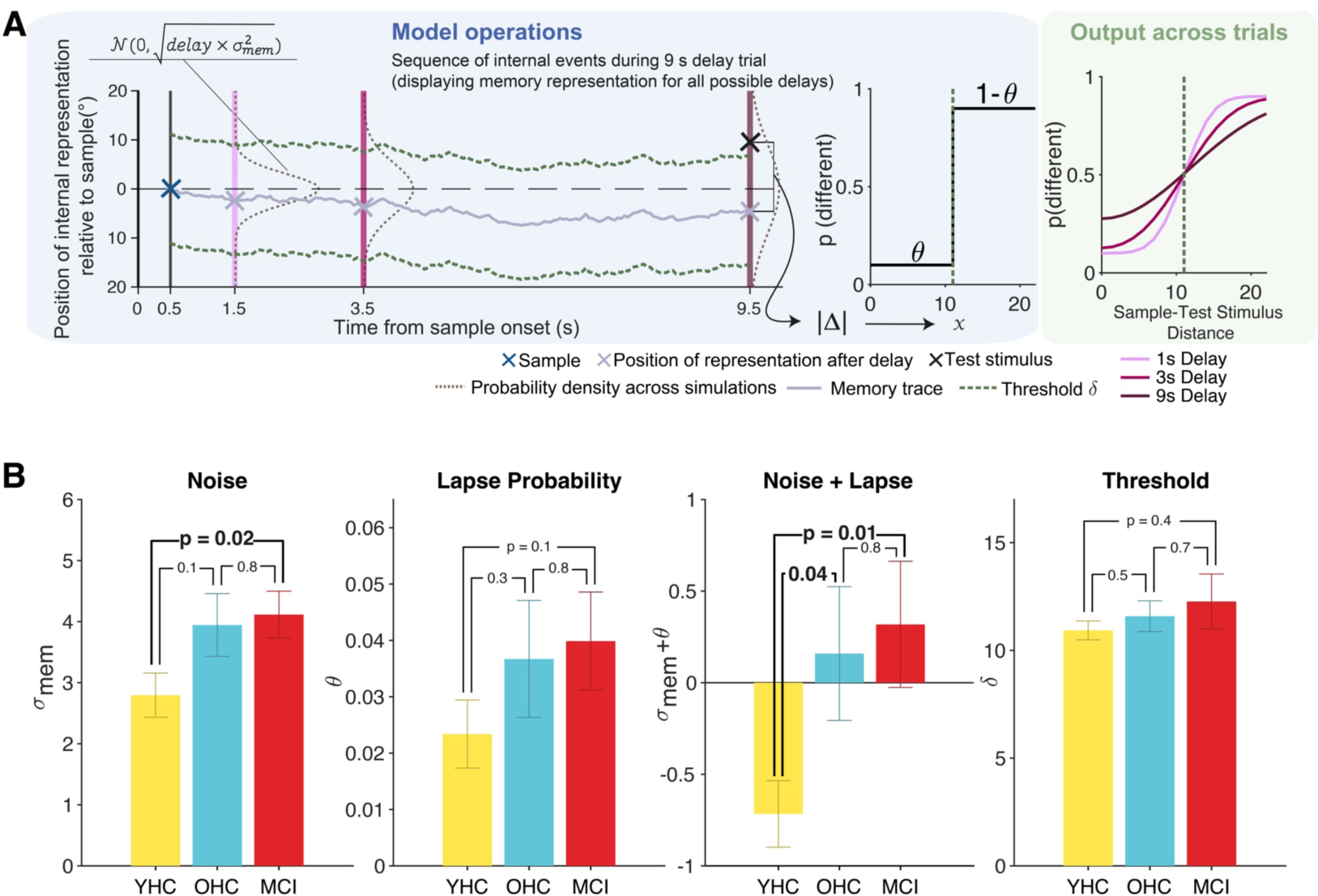
Behavioral model. **(A)** *Left:* Model schematic with one exemplary memory trace diffusing in space over time from sample (blue cross) offset (solid grey line). The diffusion causes the across-trial probability density of the memory trace to broaden over time (dashed grey lines at delay durations of 1, 3 and 9s). At the end of the delay (three purple lines corresponding to the three delay durations used here) there is a readout of the memory representation (grey crosses). In the example shown here, a “non-match” test stimulus (black cross) is presented after 9s of delay (black). The dashed green line represents the decision threshold, which produces a ‘correct’ response in this example as the test stimulus lies outside the threshold. *Middle*: Decision function (DF) for translating memory representation-test stimulus distance into probability of a “different” response, with symmetric asymptotes defined by the value of the time-independent lapse probability (θ). *Right*: Model-derived choice probabilities for reporting “different” as a function of sample-test stimulus distance for each delay duration (shades of purple). **(B)** Fitted model parameters for each subject group (mean ± s.e.m.) and *p*-values of associated between-subjects non-parametric permutation-tests.

In some model variants, we also allowed for the possibility of time-dependent memory lapses. The time-dependent memory lapse probability was modelled as a hazard function in which the likelihood of a memory lapse having occurred by time *t* during the delay (θ_*mem,t*_) accumulated over time:

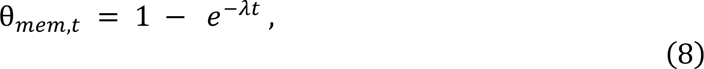

where λ was the hazard rate.

The overall choice probability was then computed as a mixture model as follows:

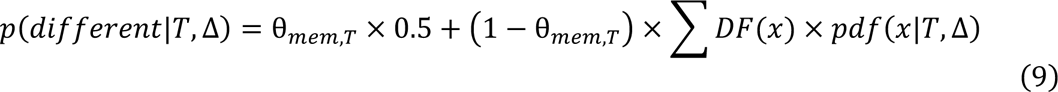

We fit six different variants of this general model to participants’ choices (Table 2).

**Table 2.** Free parameters in fitted model variants.

| Model # | Memory Noise ($\sigma_{mem}$) | Threshold ($\delta$) | Lapse ($\theta$) | Memory lapse rate ($\lambda$) | Decision noise ($\sigma_{dec}$) | # of free parameters |
| --- | --- | --- | --- | --- | --- | --- |
| 1 | X | X |  |  |  | 2 |
| 2 | X | X | X |  |  | 3 |
| 3 | X | X |  | X |  | 3 |
| 4 | X | X |  |  | X | 3 |
| 5 | X | X | X | X |  | 4 |
| 6 | X | X | X | X | X | 5 |

The most complex model variant (Model 6) allowed all five above-described parameters to vary: memory noise σ_mem_, decision noise σ_dec_, decision threshold δ, fixed lapse probability θ, and hazard rate of memory lapses λ. The simplest model variant (Model 1) fit only σ_mem_ and δ as free parameters. All variants of intermediate complexity fit σ_mem_ and δ as additional free parameters. In variants that fit only a subset of the five parameters described above, all other parameters were set to zero. Model variants not including time-independent (θ) and time-dependent (λ) lapses resulted in asymptotes of the decision function equal to 0 and 1. Model variants in which decision noise (σ_dec_) was set to zero resulted in a *DF* that took the form of a step function (Fig. 3A*, middle*) as opposed to a smooth sigmoid.

### Parameter Estimation

The objective function to be minimized during model fitting was defined as the cross-entropy across trials *trl* between the participants’ responses and model predictions for the likelihood of a “different” response:

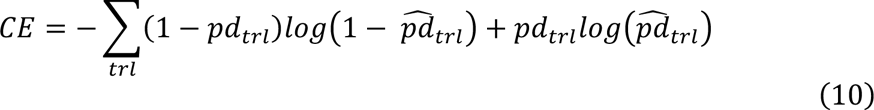

Best-fitting parameters for a particular model variant were found by minimizing this objective function using the particle swarm optimization algorithm (100 particles with wide parameter bounds and initialized at pseudorandom locations: max. of 1000 iterations) using a toolbox designed for implementation in MATLAB^38^.

### Model selection: parameter recovery and model comparison

Among the six candidate fitted models (Table 2), we selected a single variant through a combination of parameter recovery and formal model comparison, and the parameter estimates of this variant were then used for all analyses reported in ‘Results’. To evaluate the recoverability of parameters from the different candidate model variants, we simulated behavioral data sets (N=100, each consisting of 189 trials) for each variant where each parameter was set to a representative value across all subjects from fits of the least complex model that included that parameter (σ_mem_ = 4.2856, σ_dec_ = 3.0802, δ = 11.1370, θ_dec_ = 0.0203, λ = 0.0049). We evaluated recovery of σ_mem_ and δ (key parameters present in all model variants) by means of the width of the distribution of the fitted parameters across all 100 simulated datasets. This procedure showed that recoverability of σ_mem_ and δ was compromised in models that included σ_dec_ as a free parameter (Supplementary Fig. 3B), excluding those two models from further consideration.

Next, the goodness-of-fit of the remaining 4 model variants was compared by means of Bayes’ Information Criterion (BIC) scores as follows:

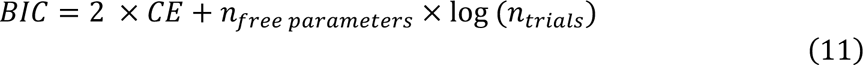

Model 2, which included σ_mem_, δ and λ as free parameters, was the model with the lowest mean BIC score when pooling all subjects across groups (Supplementary Fig. 3E). Taken together, parameter recovery analyses and model comparison motivated our selection of Model 2 (Table 2) for all further analyses of the parameter estimates (link to overt behavior, cognitive integrity, and MEG). We acknowledge that it remains possible that time-dependent memory lapses and/or decision noise affected participants’ behavior. However, our parameter recovery analysis suggests that the current experimental manipulations are not adequate for identifying the contributions of these additional parameters.

## MEG data analysis

MEG data were analyzed with a combination of customized scripts (see associated code which will be made available upon publication) and the following toolboxes: FieldTrip^39^ for MATLAB, MNE^40^ and pymeg for Python (https://github.com/DonnerLab/pymeg), as established in previous work of our laboratory^41,42^.

### Preprocessing

Preprocessing of the MEG data proceeded according to a standardized pipeline developed in our laboratory (https://github.com/DonnerLab/meg-preproc). This broadly involved initial artifact detection and removal using both independent component analysis (ICA) and non-ICA methods, trial segmentation, and reconstruction of the signal with respect to its cortical source.

Continuous MEG time series for each task block were first resampled to 400 Hz, and line noise was removed by bandstop filtering around 50, 100 and 150 Hz. Time points at which any of the following MEG artifacts occurred were then identified: head movements (translation of any fiducial coil >6 mm from the first data point in that block), muscle artifacts (*z*>20 after applying a 110–140-Hz Butterworth filter and *z*-scoring), sensor jumps (detected by Grubb’s outlier test for intercepts of lines fitted to log-power spectra computed on 7s data segments, with 20% overlap between successive segments), and other noise sources (usually due to cars passing the MEG laboratory, identified as any 2s data segment in which any sensor had data range >20 pT). The timings of blinks (Eyelink algorithm) and saccades (gaze changes >1.5 d.v.a) were identified using the eye-tracking data if these data were deemed of sufficient quality after visual inspection, or through outliers in the vertical EOG (z-score > 2). Heart beat timings were identified by applying FieldTrip’s *ft_artifact_ecg* algorithm to the ECG data.

Having detected the artifacts described above, we then high-pass filtered the continuous MEG data at 1 Hz, discarded any time points containing head movement, muscle, jump or car/other artifacts, concatenated the cleaned time-series across blocks for a given participant, and subjected the resulting concatenated data to ICA (infomax algorithm). Component time-series were then segmented from −1 to 2s around identified blinks and saccades (−0.3 to 0.3s around identified heart beats); components were ranked by their coherence with EOG (ECG) data equivalently segmented around blinks/saccades (heart beats); the temporal trajectories, spectral properties and spatial topographies of the 25 components with the highest coherences were visually inspected; and the component numbers of those judged to capture eye or cardiac artifacts were noted. In a final set of steps, the ICA weights were back-projected onto the downsampled, bandstop-filtered continuous MEG data now subjected to a high-pass filter of 0.1 Hz; those components capturing eye and cardiac artifacts were removed from the data; and the data were epoched into trial intervals from 0.6s before sample onset to 0.3s after test stimulus onset. In addition to those excluded due to the response time and accuracy criteria described above for behavioral analyses, trials with any time point containing previously-identified head movement, muscle, jump or car/other artifacts were excluded from all MEG analyses.

Subjects who despite careful recruitment were judged through visual inspection to be subject to persistent artifacts in their MEG data (presumably from presence of metallic materials, used for example in dental work), were preprocessed without discarding these time points and the data quality was re-evaluated after the source reconstruction procedure (N=7). Subjects whose data on visual inspection after source reconstruction still displayed slow fluctuations indicating metal artifacts were excluded from further MEG data analysis altogether (N=3).

Our MEG analyses focused primarily on data around sample presentation and the first second of the delay. If trials with a delay duration above one second did not contain artifacts during the first second of delay, these were preserved for the analysis together with the 1s delay trials to increase the number of trials available for analysis and computation of the data covariance matrix (see ‘Spectral analysis and source reconstruction’). If a cleaned dataset for an individual subject consisted of fewer than 80 trials after preprocessing, then that subject was excluded from all MEG analyses (N=4). In total, this procedure resulted in the exclusion of 7 of the 46 subjects who contributed to the behavioral results, leaving 39 subjects for MEG analysis in total (16 for MCI, 17 for OHC and 6 for UNC).

### Spectral analysis and source reconstruction

We first subjected the trial-averaged (phase-locked) response of each sensor from the single-trial time courses, in order to isolate activity components that are non-phase-locked to stimulus onset. The latter are generated by recurrent synaptic interactions that are also involved in the generation of persistent cortical activity^8,43^. Time-frequency representations (TFRs) of complex-valued Fourier coefficients (phase and amplitude information) for individual trials were calculated using a sliding window Fourier transform. We used Hanning tapers for the frequency range 1-35 Hz (window length: 0.4 s; time steps: 0.05 s; frequency steps: 1 Hz; frequency smoothing, ±2.5 Hz) and the multi-taper method with discrete proloid slepian tapers (window length, 0.25 s; time steps, 0.05 s; frequency steps, 4 Hz; frequency smoothing, ±6 Hz) for the frequency range 36-120 Hz.

For the source reconstruction, we used (mostly individual-subject, see below) structural MRI scans to generate three-layered head models in FieldTrip, which were in turn used to compute the forward solution (leadfield) for each source point. The cortical surface was reconstructed using Freesurfer^44,45^ and aligned to established anatomical atlases (see ‘Definition of ROIs and ROI groups’). For subjects with artifactual (N=6) or no (N=4) MRI scans, a template average surface provided by Freesurfer (‘fsaverage’) was used instead. We used linearly constrained minimum variance (LCMV) beamforming to project the sensor-level Fourier coefficients into source space, specifically: onto 4,096 vertices per hemisphere located on the cortical surface (recursively subdivided octahedron). We computed LCMV-filters with MNE using a covariance matrix of the cleaned, epoched single-trial (broadband) data to constrain a forward model. This covariance matrix was computed using the data from all trials irrespective of delay duration between 0.25s before and 1.5s after sample onset. At each vertex, the source orientation was selected based on the maximum output source power determined through singular value decomposition. To overcome random sign flips of the beamformer results, the polarity of the time series of adjacent vertices was aligned. Then the complex-value Fourier coefficients of each vertex were computed by application of the corresponding spatial filter and transformed into power by taking the absolute value and squaring.

The source-level power estimates were averaged across all vertices within each region of interest (ROI; see ‘Definition of ROIs and ROI groups’) and normalized with respect to the mean baseline spectrum using the dB transform. The baseline spectrum was computed by averaging power estimates across the interval −0.4s to −0.2s from sample onset and then across all trials. Because we had no a priori hypotheses about hemispheric lateralization effects and the anatomical atlas we used is symmetric, the power modulation values were further pooled across the left- and right-hemispheric parts of each ROI. Power modulations of the ROI groups were evaluated as the mean trial-averaged power values of the ROIs within each ROI group (see Table 3).

**Table 3.**
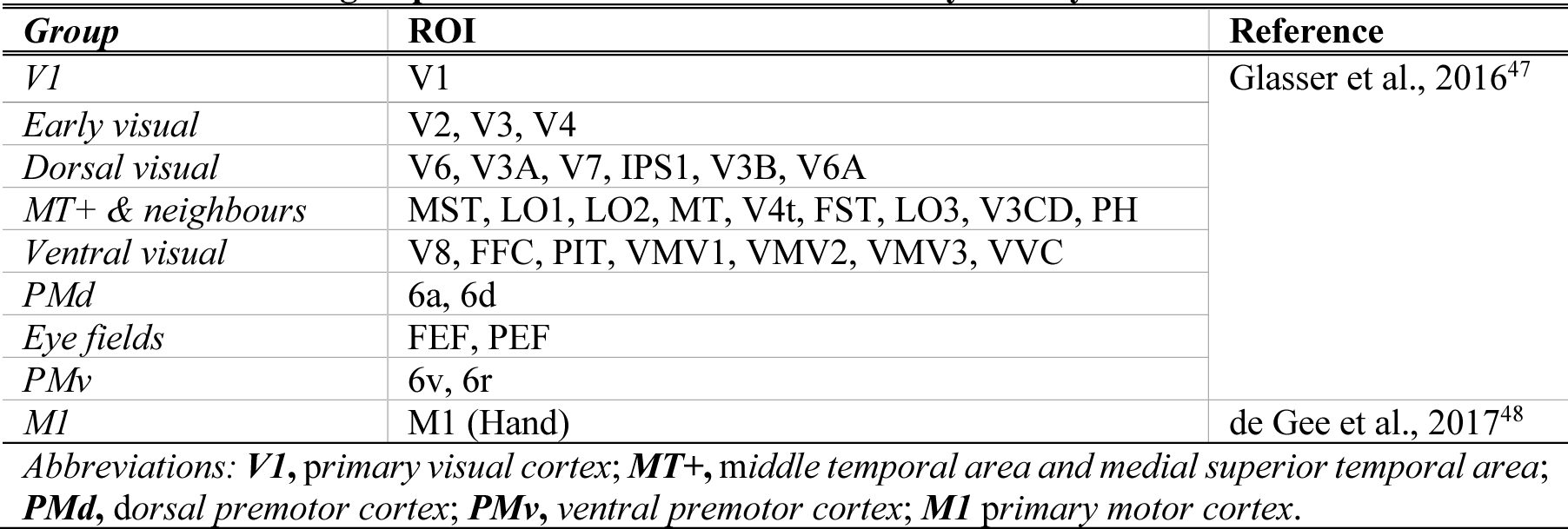
Definition of groups of ROIs for time courses of delay activity.

### Decoding of sample spatial location

We trained multivariate decoders to predict the angular position of the sample stimulus from the spatio-spectral power modulation patterns in each cortical area during the trial. Decoding was performed using ridge regression through scikit-learn for Python^46^, separately for each subject and time point. We normalized the power values of all vertices per ‘brain region’ (from both hemispheres) and 31 frequency bins (range: 5-35 Hz) by z-scoring across trials. This frequency range was chosen because we observed sustained power modulations in this range throughout delay intervals (Supplementary Fig. 5A). In separate versions of this analysis, ‘brain region’ referred to individual ROIs from the anatomical atlas or ROI groups (see Table 3), across which all vertices were pooled. To reduce the dimensionality of the data, principal component analysis was performed on the training data and we only used the components accounting for top 80% of the variance in the data for the decoding. This cutoff was chosen to avoid overfitting given the relatively low number of trials and resulted in 39.49 ± 8.5 (mean ± S.D.) components averaged across all time points (Fig. 6A) for example area V1. The resulting components were used as decoding features. The decoder was fit using 10-fold cross-validation and an L2 penalty of α = 1. Decoding precision was evaluated by Pearson correlation coefficient between the predicted sample angle and its actual angle.

### Predicting individual cognition from MEG markers of cortical delay activity

In order to relate markers of cortical delay activity to computational model-based or neuropsychological test scores, we focused on four markers derived from the spectral power modulations and evaluated across the first 1s of delay, which was available on all trials: (i) mean power modulation (*pow*) in the range 5-35 Hz; (ii) mean decoding precision (*dp*); (iii) across-trial variance (*atv*) of the time-averaged power modulation estimates; and (iv) within-trial variance (*wtv*) of power modulation estimates across 20 successive time steps within individual trials, followed by averaging variance estimates across trials. For simplicity and due to a lack of a priori hypotheses about specific key regions, we averaged these markers across all 180 ROIs (see ‘Definition of ROIs and ROI groups’) and z-scored across included subjects.

We then fit different multiple linear regression models to predict different individual cognitive measures (model parameter estimates as well as cognitive integrity scores) from these four cortex-wide neural markers of delay activity:

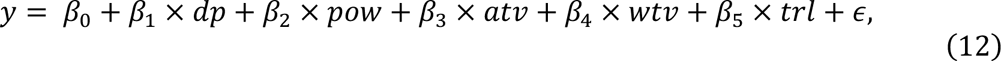

where *y* were, in different model fits, parameter estimates from the behavioral model or individual cognitive integrity scores, and *trl* were individual trial counts (z-scored) that were included as nuisance regressor to account for inter-individual differences in the trial counts available for MEG analyses. This analysis was performed including all subjects in a single model, as well as for OHC and MCI groups separately.

### Definition of ROIs and ROI groups

We based the anatomical definition of ROIs on a multimodal MRI-based parcellation of the cerebral cortex (180 ROIs)^47^. For certain analyses focusing on specific regions along the visuo-motor cortical pathway, we collapsed results across multiple ROIs belonging to certain ROI groups (defined in the Supplement of Glasser et al., 2016^47^) as described in Table 3.

Here, only the primary motor region was defined independently – on the basis of previous work of our laboratory – as the hand-specific area^48^. For calculating whole-cortex maps of certain statistics, we used the 22 ROI groups defined in the supplementary text of Glasser et al., 2016^47^.

### Statistical testing

Within-subjects tests were non-parametric permutation tests against zero (10,000 permutations). For comparisons between groups of subjects, we used between-subjects non-parametric permutation tests (10,000 permutations).

Deviations from zero in ROI-or ROI-group-specific time-frequency representations and decoding time courses (Fig. 5A, Fig. 6A) were identified through cluster-based permutation testing (10,000 permutations, cluster-forming threshold *p*<0.05). Maps of measures plotted over the entire cortex (brain maps) were thresholded through FDR-correction at a significance threshold of *p*<0.05.

All correlation analyses were performed using Pearson correlation (two-tailed).

### Data availability

Analysis code will be made available upon acceptance. For reasons of patient data protection raw behavioral data and preprocessed MEG data will be made available upon reasonable request.

## Results

We used an integrative approach to gain insight into aging effects on the stability of working memory representations, and to relate these to clinical measures of cognitive integrity (Fig. 1A). Through behavioral modeling, we quantified working memory dynamics in terms of latent variables. For all older participants, we measured their cortical population activity using MEG, and their performance in a neuropsychological test battery. We combined categorical and dimensional approaches in our analysis of working memory mechanisms by (i) comparing model- and MEG-based measures between groups (OHC, MCI, and – as a reference for behavioral modeling – YHC); and (ii) for the older participants, correlating our measures with a clinical assessment of their cognitive integrity.

The first principal component of the scores from the CERAD-Plus test battery^29–31^ consisted of positive contributions of performance scores in all tests (Fig. 1B, *left*, inset) and explained 42.5% of variance in the test data (Fig. 1B, *left*). The eigenvalues (scores) associated with this component were strongly (i) correlated to alternative summary metrics (CERAD ‘total score’^34^, Fig. 1B, *middle*; and Mini Mental State Examination^49,50^, Supplementary Fig. 1B) and (ii) predictive of the clinical MCI diagnosis (ROC value ≈ 0.86, Fig. 1B, *right*). In the following, we used these scores (denoted as “PC1 score”) as a summary measure to quantify each older individual’s cognitive integrity.

### Impaired working memory performance in older adults

In our delayed-match-to-sample working memory task, subjects judged whether a ‘sample’ and a ‘test’ stimulus had the same or a different spatial location (Fig. 2A). As expected from previous work ^51–53^, the difficulty of this judgment depended on both the temporal and spatial distance between sample and test (Fig. 2B-C, Supplementary Fig. 2B-C), with the error rate increasing with delay duration (collapsing across match and non-match trials; main effect of delay in mixed delay*group ANOVA: *F_2,126_* = 38.6, *p*<10^−4^, Fig. 2B) as well as with test-sample distance (non-match trials only; main effect of distance in mixed distance*delay*group ANOVA: *F_1,63_* = 258.84, *p*<10^−4^, Supplementary Fig. 2C).

Comparison of error rates between the different participant samples (YHC, OHC, and MCI) revealed a main effect of group (mixed delay*group ANOVA; *F_2,57_* = 3.81, *p* = 0.028) and interaction between group and delay (*F_4,114_* = 2.78, *p* = 0.030). Previous modeling of spatial delayed match-to-sample performance showed that false alarms on trials with small test-sample distance are particularly informative about the stability of working memory representations^23^. Indeed, while we observed no effects of group on misses and false alarms for ‘far’ trials (Supplementary Fig. 3B-C), younger healthy controls performed better than older adults on the near false alarm trials, for both 1s and 3s (but not 9s) delays (Fig. 2C, *left* and *middle*; planned comparisons, two-sided permutation tests). There was a similar, but only trending effect for an increased task accuracy of OHC compared to MCI on 9s near trials (Fig. 2C, *right*).

### Model-based dissection of working memory performance

The behavioral effects reported above may, in principle, be due to changes in the stability of the working memory representation or in the response strategy of the subjects. Both factors may change with age. To disentangle these possibilities and to gain deeper mechanistic insight, we developed a model of working memory dynamics inspired by previous cortical circuit modeling work^20,23,37^. These biophysically detailed models consist of many parameters, which precludes fitting them to behavioral data in a principled fashion. To estimate individual parameters quantifying the mechanisms governing working memory performance, we therefore opted for a more abstract (‘algorithmic’) modeling approach using a small set of free parameters that were sufficiently constrained by our behavioral data.

Our approach assumed that instability of a working memory representation may originate from two sources. First, random drift of the activity pattern in the neuronal population encoding the sample stimulus (i.e., spatial location) will introduce random error in the information encoded at the end relative to the start of the delay period.^23,54,55^ This was captured by describing the sample representation during delay as a particle subject to a random diffusion process (Fig. 3A, *left*)^25,26^ with a diffusion constant that captured the memory noise. Second, the activity pattern may vanish altogether before the end of the delay period.^53^ In the simplest model variant tested, this was captured by a single lapse parameter (see below and ‘Methods’). For the same/different judgment required by our task, the model computed the absolute distance between memory representation at the end of delay and the then-shown test stimulus, resulting in the decision variable. The judgment was then produced through the application of a threshold (hard cutoff or smooth function; see below) to this decision variable (Fig. 3A, *middle*).

We fit the behavioral data with a selection of model variants differing in the composition of free parameters (see ‘Methods’). Model validation and comparison (Table 2; Supplementary Fig. 3A-C) favored a model variant containing the following free parameters: memory noise (σ_mem_), decision threshold (δ) and lapse probability (θ). Here, the decision translating the difference between the memory representation and test stimulus locations into a same/different response was modelled as a deterministic process (i.e., step function). The probability of lapses (i.e., random responses) was reflected in symmetric asymptotes of this function (Fig. 3A, *middle*). The model yielded probabilities of “different” judgments as a function of delay and sample-test stimulus distance (Fig. 3A, *right*). We used this model variant for all analyses described below.

The predictions of the fitted model were largely consistent with participants’ performance for all participant groups and conditions assessed here (Fig. 2B-C, Supplementary Fig. 2B-C). Further, we found expected correlations of the fitted model parameters with model-free metrics of behavioral performance based on signal detection theory: threshold δ was positively correlated with signal detection theoretic criterion *c* (Supplementary Fig. 3F, *left*) and both memory noise (σ_mem_) and lapse (θ) parameters were negatively correlated with sensitivity *d’* (Supplementary Fig. 3F, *middle, right*; all Pearson correlations, *p*<10^−4^).

The lapse parameter in this model variant captured lapses occurring at different levels of processing (sensory encoding, memory maintenance, decision, and action selection), which were not dissociable in the current task. We further note that, while the decision function is certainly an oversimplification, model comparison did not favor a model with a noisy decision transformation (Supplementary Fig. 3A-C), likely because the data also did not allow for disentangling behavioral variability due to lapses versus decision noise.

The model revealed smaller memory noise (σ_mem_) for young healthy adults compared to MCI patients and a trending effect compared to the older healthy controls (Fig. 3B, *left*). We combined memory noise and lapse rate into a single measure that captured all stochasticity in internal processing distinct from strategic sources of error (i.e., decision threshold δ). This revealed larger behavioral stochasticity in both groups of older adults than the younger healthy controls (Fig. 3B, *third from left).* There was no evidence for group differences in other model parameters (decision threshold, lapse probability on its own; Fig. 3B).

We also observed that the parameter estimates for σ_mem_ were correlated, across subjects, to those for δ (Pearson correlation, *r* = 0.25, *p* = 0.04; Supplementary Fig. 3D). We reasoned that this correlation may be genuine, reflecting a strategic adaptation to participants’ individual levels of memory noise, applying higher thresholds for “different” decisions (i.e. higher values for δ) for higher memory noise (increased σ_mem_). To evaluate the plausibility of this idea, we assessed whether such strategic threshold adjustment would help to maximize performance under larger levels of σ_mem_. Through simulation (three task blocks = 189 trials) we computed the δ settings that lead to highest accuracy for 100 different levels of σ_mem_ covering the range of fitted noise parameters observed in the subjects of the study. The performance-maximizing δ values were identified using simplex search minimizing the objective: 1 − *P*(*correct*), where *P*(*correct*) is the average response accuracy across trials. Indeed, a more conservative threshold (higher δ) produced better task performance when memory noise was high (Pearson correlation, *p*<10^−4^; Supplementary Fig. 3E).

In sum, the group differences in behavioral performance of the working memory task were due to age-related deterioration of the stability of the working memory representation, rather than (potentially compensatory) strategic changes in decision thresholds.

### Relating working memory mechanisms to cognitive integrity

We next used a dimensional approach focusing on individual differences to link the working memory mechanisms to cognitive integrity. Cognitive aging is characterized by substantial differences between individuals, which are only partly captured by the clinical classification into older healthy controls and MCI categories (Fig. 1). For example, seven older participants who were originally recruited for our OHC group exhibited neuropsychological test scores comparable with MCI and were, therefore, classified as a separate cohort for the purpose of this study (UNC, Fig. 1A). In the following, we used each participant’s neuropsychological test results, summarized by the eigenvalue of the first principal component (PC1, Fig. 1B), as an individual measure of overall cognitive integrity.

Individual cognitive integrity was robustly correlated to performance on the working memory task as well as the model parameters capturing behavioral stochasticity (in particular: stability of memory representation): Higher cognitive integrity was associated with higher task accuracy and lower combined memory noise/lapse scores from the model (Fig. 4A-B). Remarkably, the correlation with task accuracy was present only in the MCI but not the OHC group individually, with a clear difference in correlation between the two groups (Fig, 4A). Likewise, there was a correlation to cognitive integrity for noise and lapse rate combined in the MCI but not the OHC group, with a trend toward a difference in correlations between the groups (Fig. 4B). This correlation in the MCI group was primarily driven by memory noise over lapse rate (Supplementary Fig. 4A). By contrast, we observed no correlation between cognitive integrity and the decision threshold parameter (Fig. 4C).

**Figure 4.**
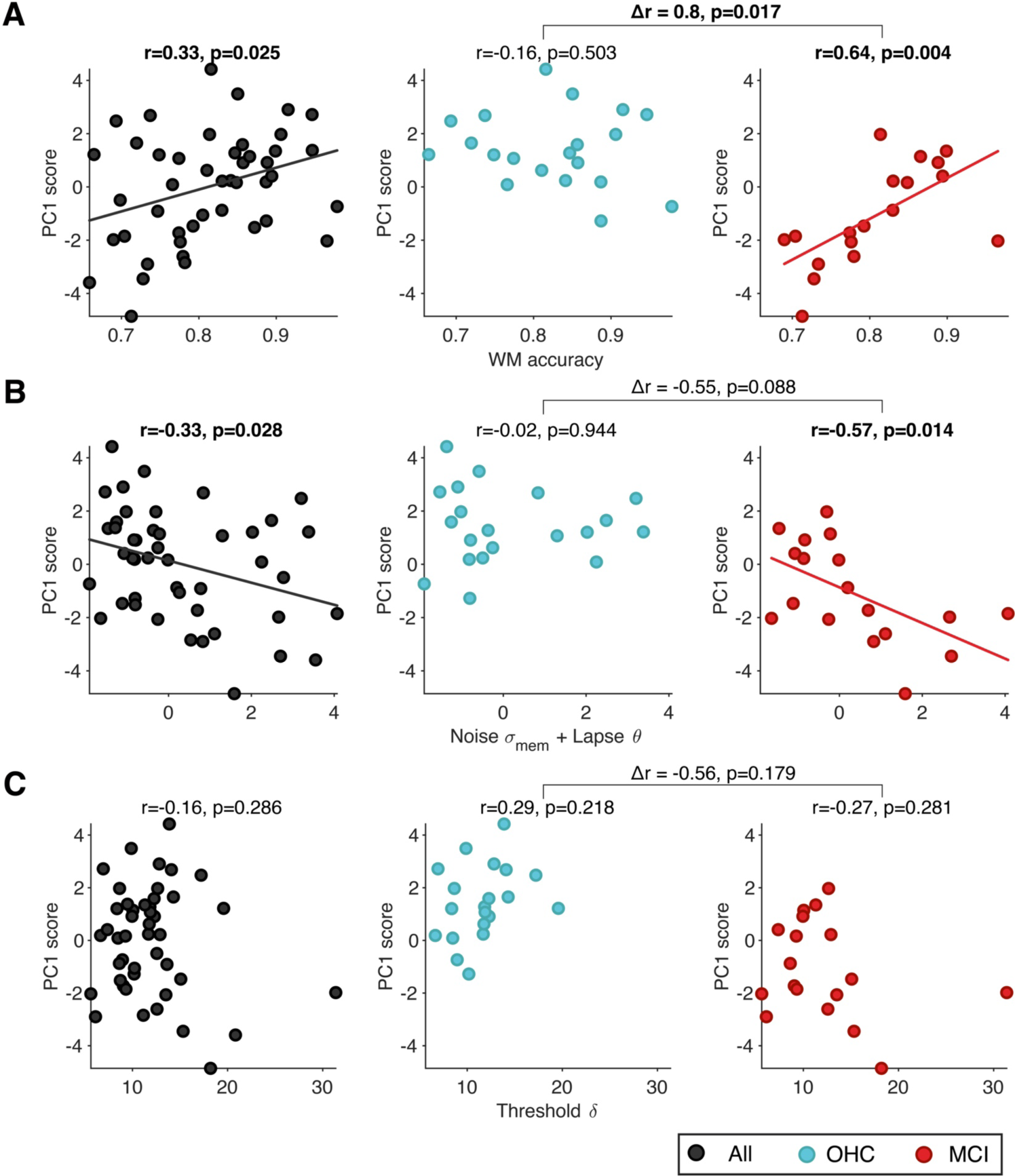
Correlation of cognitive integrity and working memory mechanisms. Correlation of PC1 scores (quantifying overall cognitive integrity, see Figure 1B) with **(A)** working memory task accuracy, **(B)** combination of memory noise and lapse probability fits, and **(C)** threshold fits. In all panels, circles represent individual subjects within the group of older participants. Correlations and correlational statistics are reported for all older subjects (N=45, black, *left*), OHC (N=20, blue, *middle*) and MCI (N=18, red, *right*) separately. Linear regression fit is shown for statistically significant correlations only. Differences in correlations between OHC and MCI are shown on top of the square brackets, associated p-values refer to two-sided permutation tests.

In sum, we established a link between individual cognitive integrity on the one hand and working memory task performance as well as model-estimated memory noise parameters on the other hand, specifically in the MCI group. Our final analyses aimed to relate the modeling results to direct measurements of cortical delay activity underlying the working memory task.

### Task-related cortical dynamics and relation to model parameters

The combination of spectral and source analysis with anatomical atlases^41,42^ yielded a detailed description of the task-related cortical population dynamics (Fig. 5). The test stimulus elicited an expected pattern of MEG power modulations^43^: a transient increase in the <8 Hz and 50-100 Hz (gamma) frequency ranges in visual cortical areas, followed by suppression in the 8-36 Hz (alpha/beta) frequency range, which was sustained throughout the delay interval (Fig. 5A, Supplementary Fig. 5A) and widely distributed across posterior cortex (Fig. 5B). The magnitude and spatial distribution of this power suppression during delay was similar in the OHC and MCI groups, without any evidence for differences (Supplementary Fig. 5C).

**Figure 5.**
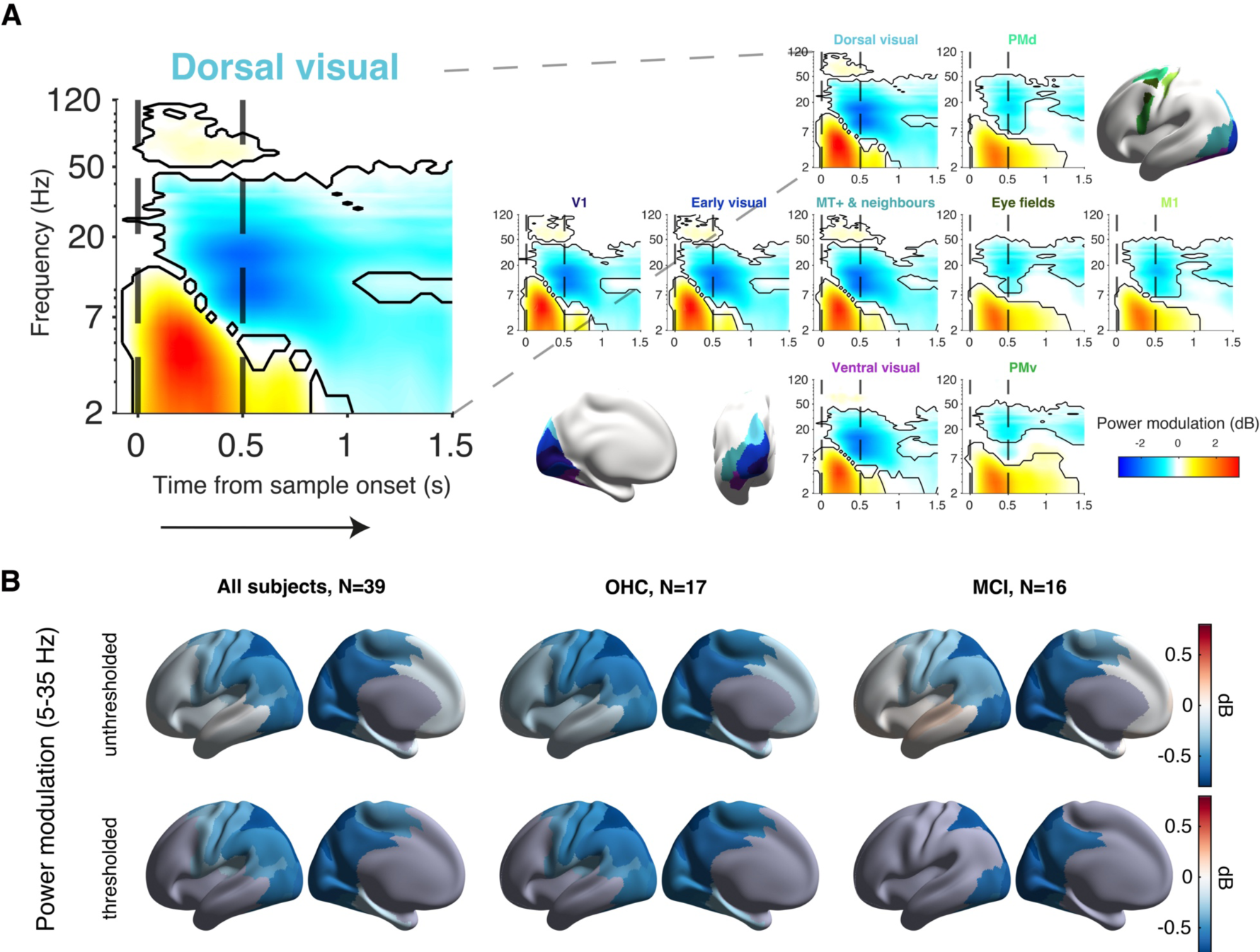
Cortical population dynamics during working memory task. **(A)** Time-frequency representations of task-induced power modulations across cortical visuo-motor pathway. Data are shown from 0.1s before sample onset (first vertical black dashed line) until 1s after sample offset (second black dashed line), and collapsed across all older adults (N=39). Black contouring captures time-frequency clusters of significant within-subject power modulations (*p*<0.05, cluster-based permutation test). Brain maps at inset depict clustered brain regions of interest (see also Table 2), colored to match panel titles of heat maps (A). **(B)** Maps of low-frequency power modulation (averaged over 5-35 Hz and 0.5s – 1.5s). *Left to right:* all older participants (N=39), OHC (N=17), MCI (N=16). *Top*: Not thresholded; *bottom*: statistically thresholded difference maps (*p*<0.05, FDR-corrected).

We used spatio-spectral features (Methods and ref. ^42^) to decode the spatial location of the sample in the local cortical population activity (Fig. 6A). While the sensory response produced the highest decoding precision during sample presentation, all visual cortical areas continued to show above-chance decoding during delay, most pronounced and sustained in dorsal visual cortex (Fig. 6A-B, Supplementary Fig. 5D). Like the power modulations, decoding precisions were similar for both groups (Fig. 6B, Supplementary Fig. 5D).

**Figure 6.**
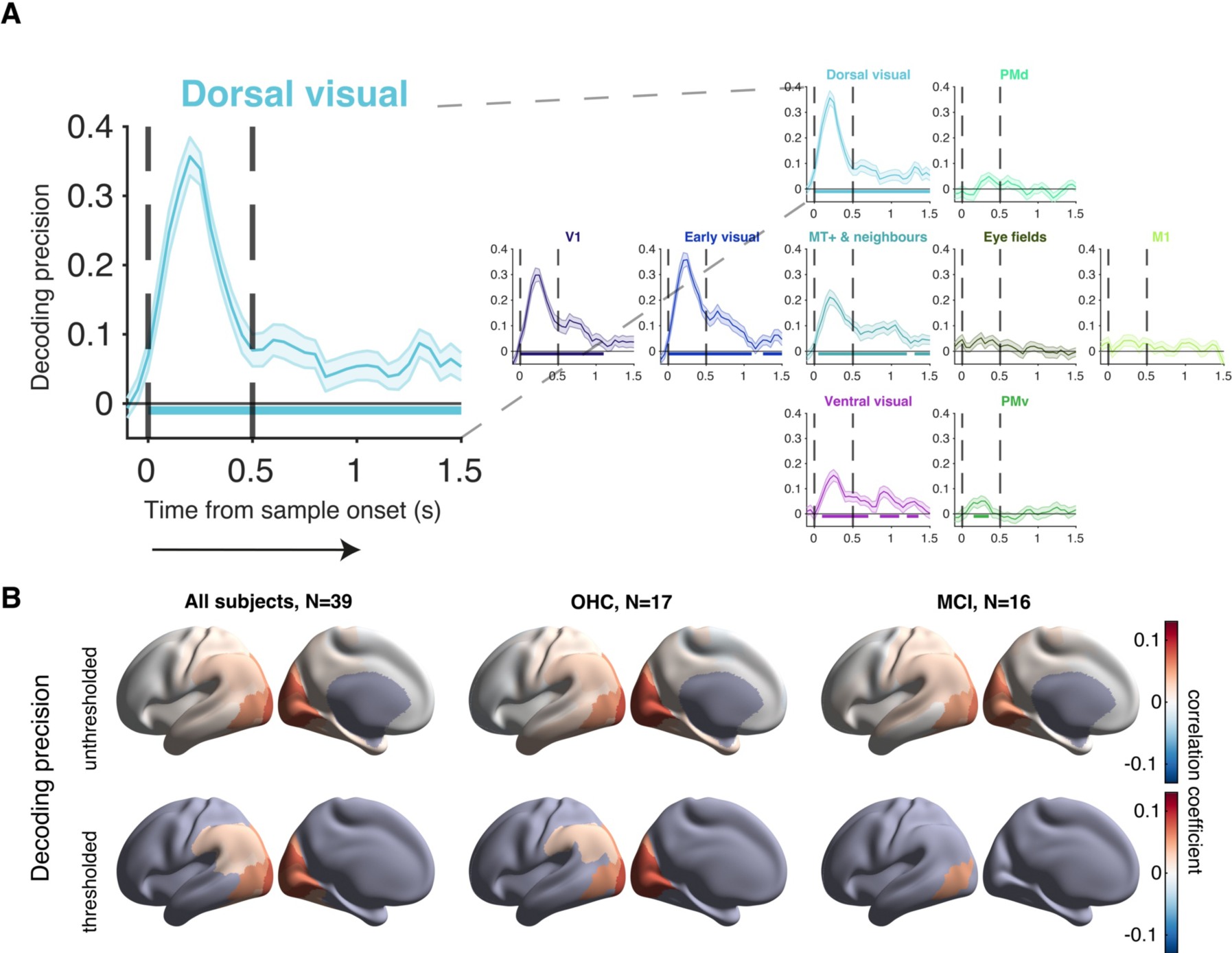
Cortical dynamics during working memory task. **(A)** Time courses of vertex-based sample location decoding precision (see text) for clustered brain regions across the visuo-motor pathway (all older subjects, N=39; mean ± s.e.m.). Times of sample onset and offset are marked by vertical black dashed lines. Colored horizontal lines indicate latencies of significant decoding precision (p<0.05, cluster-based permutation test). **(B)** As Fig. 5B, but for decoding precision during delay (0.5s – 1.5s).

Our final set of analyses related the behavioral modeling parameters to different markers of cortical delay activity during our task. We quantified four measures of the delay activity (from 0.5 to 1.5 s): in addition to mean power modulation (5-35 Hz) and decoding precision (as shown in Figs. 5 and 6), also the within- and across-trial variability of these power modulations. Measures of neural variability were included because neural variability has been implicated in cognitive integrity and aging.^2,56–58^ Due to the widespread distribution of decoding precision and overall power modulation, we collapsed each measure of delay activity across all cortical ROIs (N=180). We then fit a series of linear models regressing individual cognitive performance measures onto these activity markers (alongside some nuisance regressors).

These neural markers were not linked to individual cognitive integrity (PC1 score; Supplementary Fig. 6A-B; all *p*>0.05). But higher neural decoding precision predicted higher working memory task accuracy across subjects (similar trends within each group; Fig. 7A). None of the three other neural measures showed such a relationship. This effect was explained by the link between decoding precision and behavioral stochasticity parameters (memory noise and lapse combined; Fig. 7B), not decision threshold (Fig. 7C), again with a similar trend within each group (significant for MCI, but not OHC).

**Figure 7.**
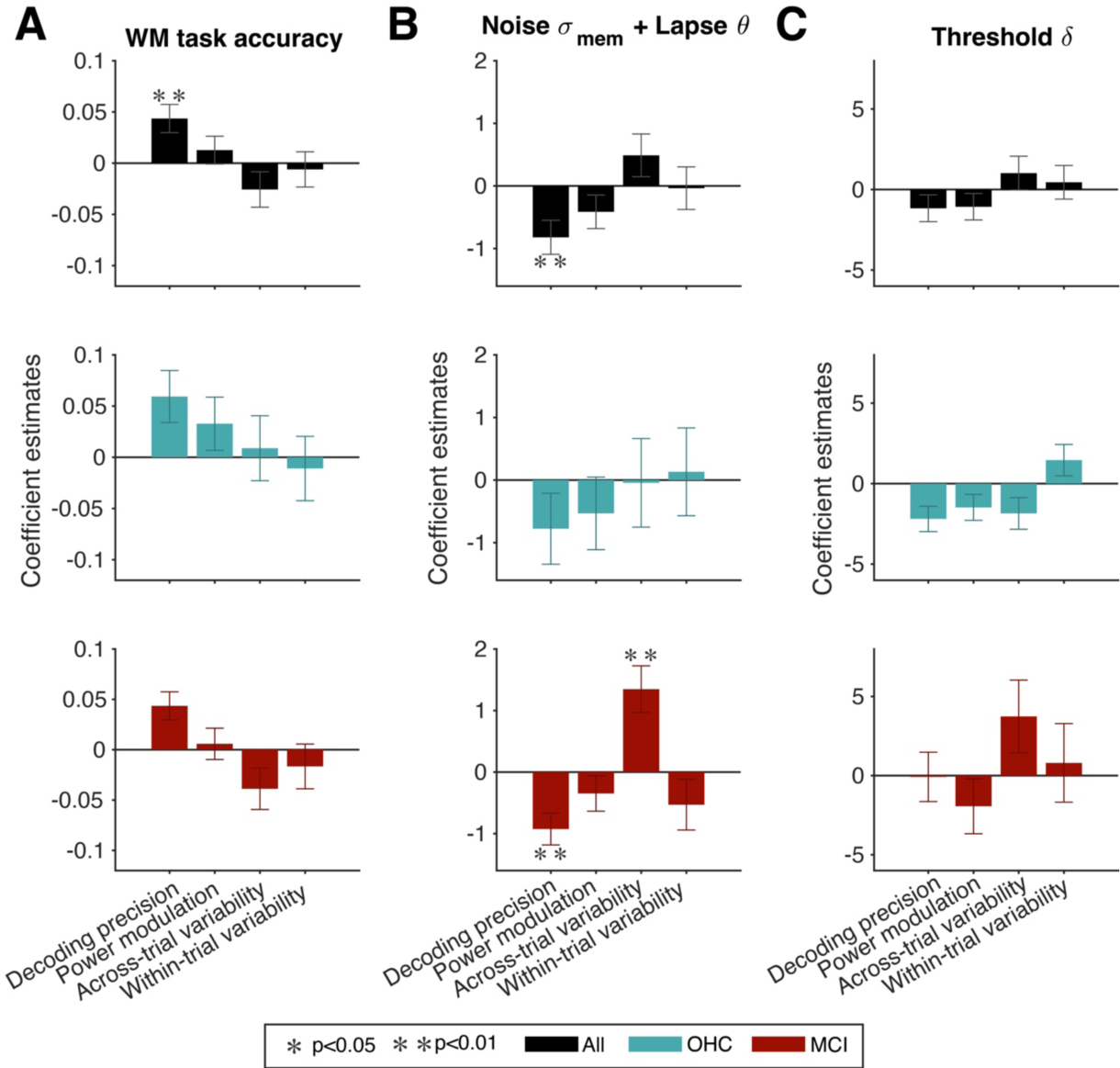
Linking markers of cortical delay activity to working memory stability. Coefficient estimates (± standard error) from multiple linear regression models predicting **(A)** working memory accuracy, **(B)** the combination of memory noise and lapse parameters, and **(C)** threshold across older individuals (*black*), OHC (*blue*) and MCI (*red*) using standardized versions of the four neural measures (x-axis) evaluated across the entire cortex. *, p<0.05; **, p<0.01 (FDR-corrected, non-significant otherwise).

Notably, the analysis also revealed across-trial variability of power modulations as a second significant predictor of individual differences in the behavioral stochasticity parameters (memory noise and lapse probability combined), specifically in the MCI group (Fig. 7B), and again with no effect for threshold (Fig. 7C).

Taken together, our MEG results support the notion that the behavioral model-derived parameters are useful markers of the neural mechanism of working memory and point to the potential relevance of trial-to-trial variability of power modulations as a marker of MCI.

## Discussion

Age-related changes in working memory performance have been investigated using a variety of task protocols. This previous work has revealed diminished working memory capacity with age in contexts requiring maintenance of multiple memoranda^e.g. 59^, especially when conjunctions of different object features (rather than single features) needed to be remembered.^60^, but see^61^ Less is known about the quality of working memory representations in human aging. One landmark study of aging monkeys has identified a deterioration of stimulus-selective persistent activity in prefrontal cortex.^13^ It has remained unknown whether and how these neurophysiological changes generalize to the human brain, how they relate to working memory performance, and to individual cognitive integrity. Our current study provides a comprehensive assessment of age-related changes in the stability of performance-relevant working memory representations, their neurophysiological bases, as well their links to individual cognitive integrity in a clinically well-characterized sample of older participants.

One of the main insights afforded by our model-based approach is that reductions in working memory performance with aging (compared to younger adults) are due to a deterioration of the stability (quality) of working memory representations, rather than a change of task strategy. The latter was captured by our threshold parameter, which governed how liberal or conservative subjects were in their (same vs. different) judgments. The model-inferred working memory quality, in turn, related to individual differences in cognitive integrity within the group of older adults, establishing its broad functional relevance. These conclusions invite comparison with studies of age-related changes in perceptual decision-making.^62^ Some behavioral modeling work has revealed that the increases in reaction times in two-choice tasks commonly observed in older adults are not due to less efficient integration of decision evidence, but to increased response caution, a strategic effect.^62^ Given the close mechanistic links between working memory and evidence accumulation in deliberative decisions^26,64^, one might expect that the increase in memory noise we inferred in older adults will translate into less efficient evidence accumulation (i.e., lower drift rate)^62^, at least across longer timescales at which mechanisms of persistent activity play out. Indeed, recent work constraining behavioral model fits with neurophysiological data provided first evidence for degraded evidence accumulation in aging.^63^ It will be instructive to study aging effects in evidence accumulation tasks that are more protracted^e.g. 41,65,66^ and/or more cognitive in nature^67,68^ than the ones used in the previous aging literature.

The relationship between our behavioral markers of working memory quality (noise and lapse rate) and overall cognitive integrity was particularly pronounced in older individuals diagnosed with MCI and not in age-matched healthy controls. This indicates a tight link between deterioration of working memory mechanisms on the one hand and pathophysiological aging as gauged by standard comprehensive neuropsychological test batteries on the other hand. Likewise, the relationship between MEG markers of cortical delay activity measured during the delay, especially for trial-to-trial neural variability, and the above model-derived markers (memory noise and lapse rate) was also particularly strong in the MCI group. Taken together, these observations highlight the translational potential of model- and MEG-based assessments of working memory mechanisms for the objective detection and monitoring of cognitive impairment during aging.

Our MEG approach builds on recent neuroimaging work that illuminated the possibility of decoding the content and precision of stimulus-selective working memory representations from non-invasively measured spatial patterns of brain activity. Previous functional MRI studies have decoded stimulus-specific information from activity patterns in frontal, parietal and visual cortex during working memory delays.^15,16^ While similar decoding approaches to event-related potential signals failed to identify lasting stimulus-selective activity patterns^69–71^, they commonly succeeded when applied to modulations of band-limited signal power^17–19^, as we did here. These stimulus-selective activity patterns exhibit hallmarks of persistent activity states^18^. Moreover, while the decodability of memoranda from these signals may decrease to some degree over memory delays, this can be explained at least in part by the drift in the underlying memory representations that is thought to be a primary source of error in behavioral working memory reports.^55,72^ These characteristics – some of which we observed in patterns of spectral power modulation in dorsal visual cortex – are consistent with cortical circuit models in which working memory emerges from sustained attractor states that drift over time.^20,23,54,55^

Our results complement other studies linking behavioral performance^23^ and neural signals^17,18^ in working memory tasks to the dynamics of cortical circuit models. These links formalize relationships between neural micro-circuit properties (e.g., the ratio of recurrent excitation and inhibition in a circuit, E/I) and patterns of task behavior. Thus, such links open opportunities for understanding behavioral effects of cognitive aging in terms of underlying neural circuit properties. For example, there is indirect evidence for alterations of cortical E/I in Alzheimer’s disease.^73,74^ One challenge associated with cortical circuit modeling of measured behavioral changes is that the parameter space of cortical circuit models is high-dimensional, and in some cases different circuit alterations can produce highly similar behavioral effects.^75^ Here, we circumvented this issue by fitting a lower-dimensional model, which was well-constrained by the behavioral task while exhibiting straightforward relationships to more detailed circuit models.^23^ Complementing such behavioral modeling with non-invasive measures of E/I^76–79^ and other circuit properties may help to add much-needed constraint in future applications of these neural circuit models to empirical data.

## Acknowledgements

G.M. acknowledges the support from the Rosa-Luxemburg-Stiftung. Further we thank Emma Krink and the other student assistants, who supported the data collection of this study.

## Funding

This work was supported by the following sources: Deutsche Forschungsgemeinschaft (DFG), projects DO1240-3-1, DO1240_4-1, and SFB936/A7 (all to T.H.D.). EU Horizon 2020 research and innovation programme, Marie Skłodowska-Curie grant agreement No. 843158 (to P.R.M.). Efficacy of a therapeutic videogame in patients with mild cognitive impairment. BGV Hamburg, Z79004 2017-2024, G500-02.10/10,011 (to J.G.).

## Competing interests

The authors report no competing interests.

## Supplementary figures

**Supplementary Figure 1.**
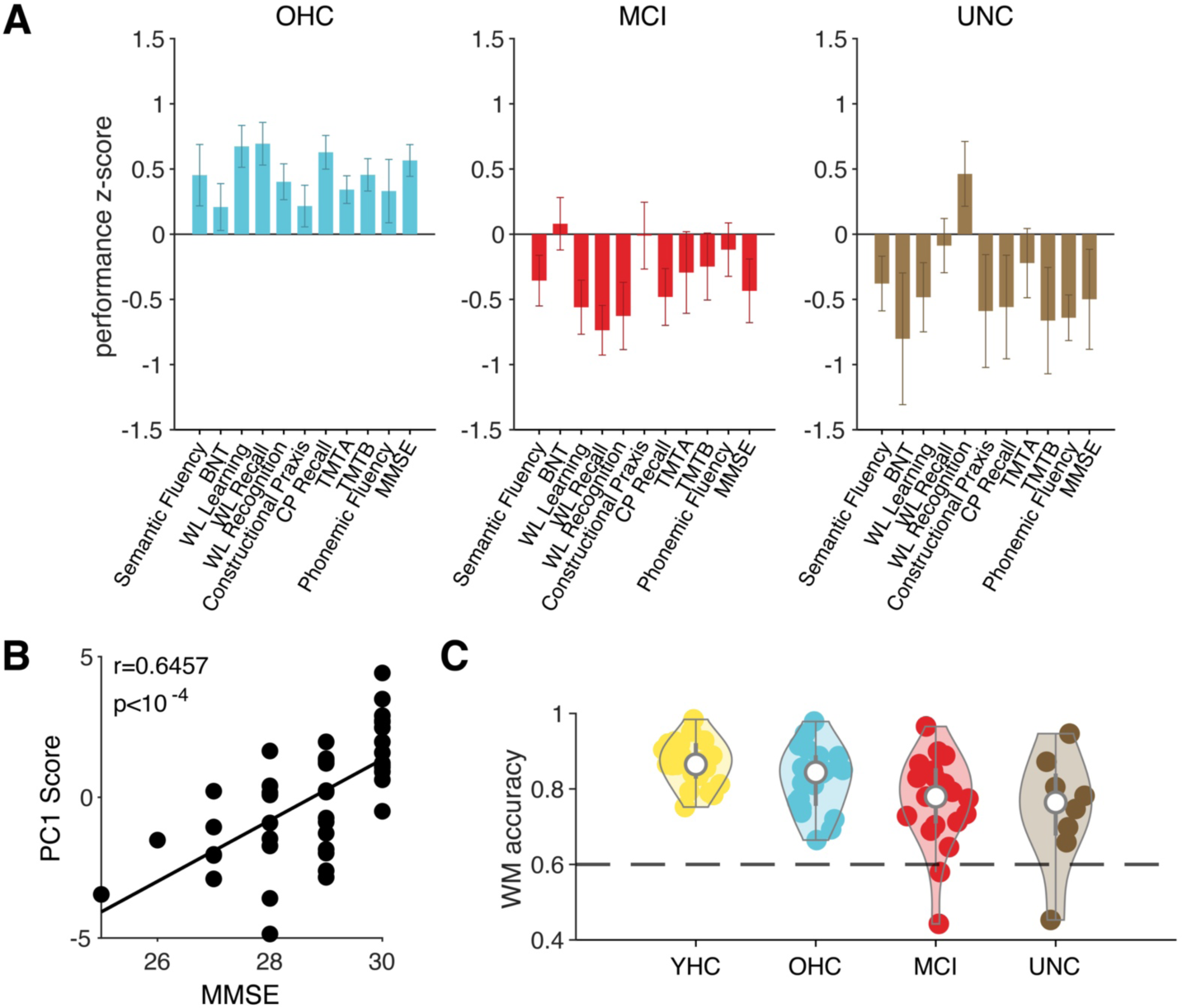
CERAD+ test battery performance scores, validation of composite cognitive score and performance exclusion criterion. **(A)** Z-scored CERAD-Plus test results (input to principal component analysis) per group (mean ± s.e.m.) show cognitive performance decline in the MCI and UNC groups compared with the OHC group. **(B)** PC1 scores and results of Mini-Mental State Examination are strongly correlated. **(C)** Single subject working memory task accuracy and kernel density estimates of subgroups show the distributions of performance measures. The application of task accuracy exclusion criterion at p(correct) = 0.6 (dashed black line) leads to the exclusion of three subjects (MCI, N=2; UNC, N=1).

**Supplementary Figure 2.**
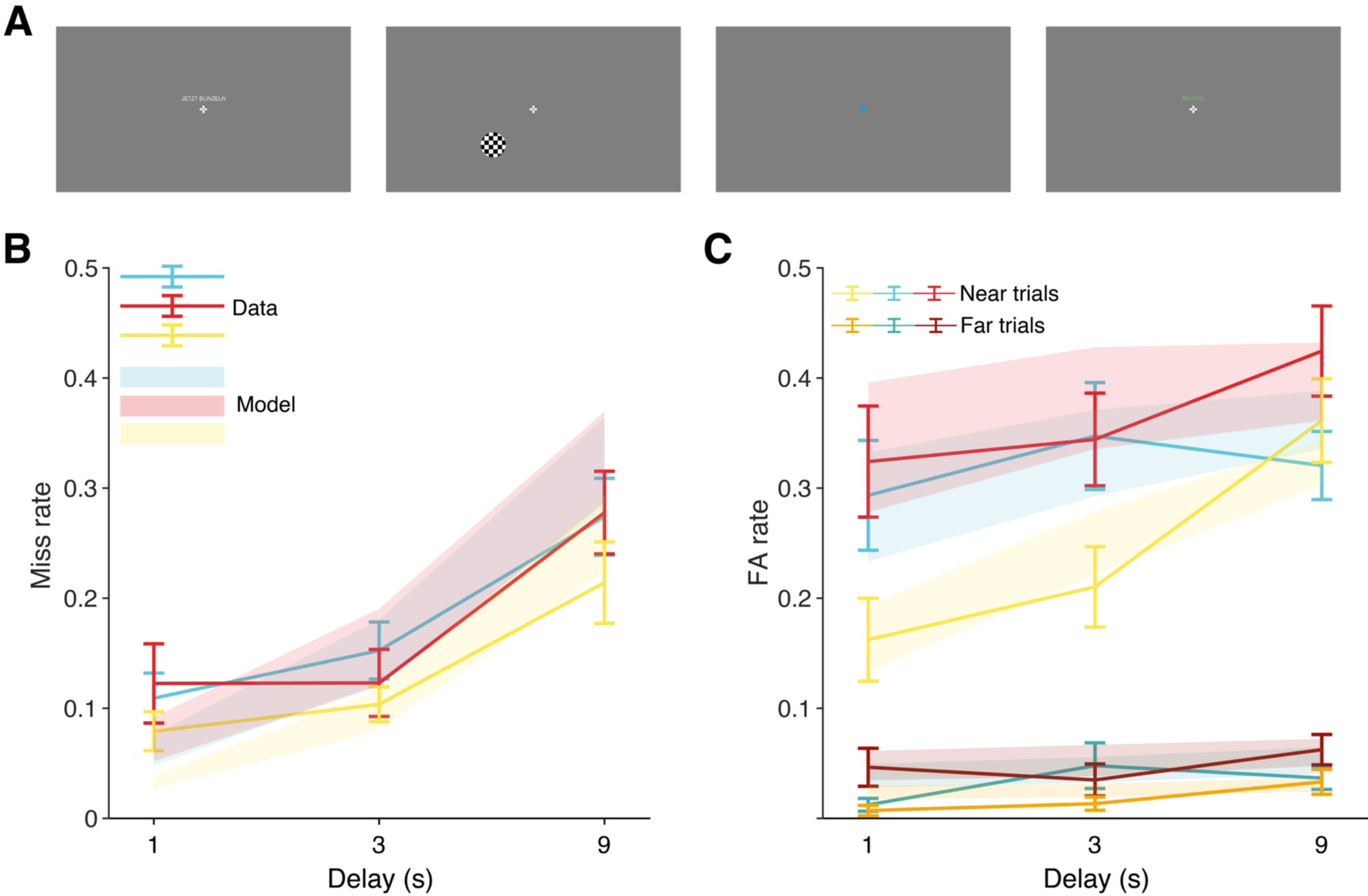
Original task frames, and error rates on working memory task differentiated by delay duration, sample-test stimulus distance, and participant group. **(A)** True-sized representation of the stimuli shown to the subjects. *Left to right:* break cue, fixation cross and checkerboard patch, response cue, feedback (correct). **(B)** Miss rates (mean ± s.e.m.) by delay duration for YHC (yellow, N=21), OHC (blue, N=20) and MCI (red, N=19). **(C)** False alarm rates (mean ± s.e.m.) on near non-match trials (lighter shades) and far nonmatch trials (darker shades). Model predictions (mean ± s.e.m.) are shown as shadings in the corresponding color.

**Supplementary Figure 3.**
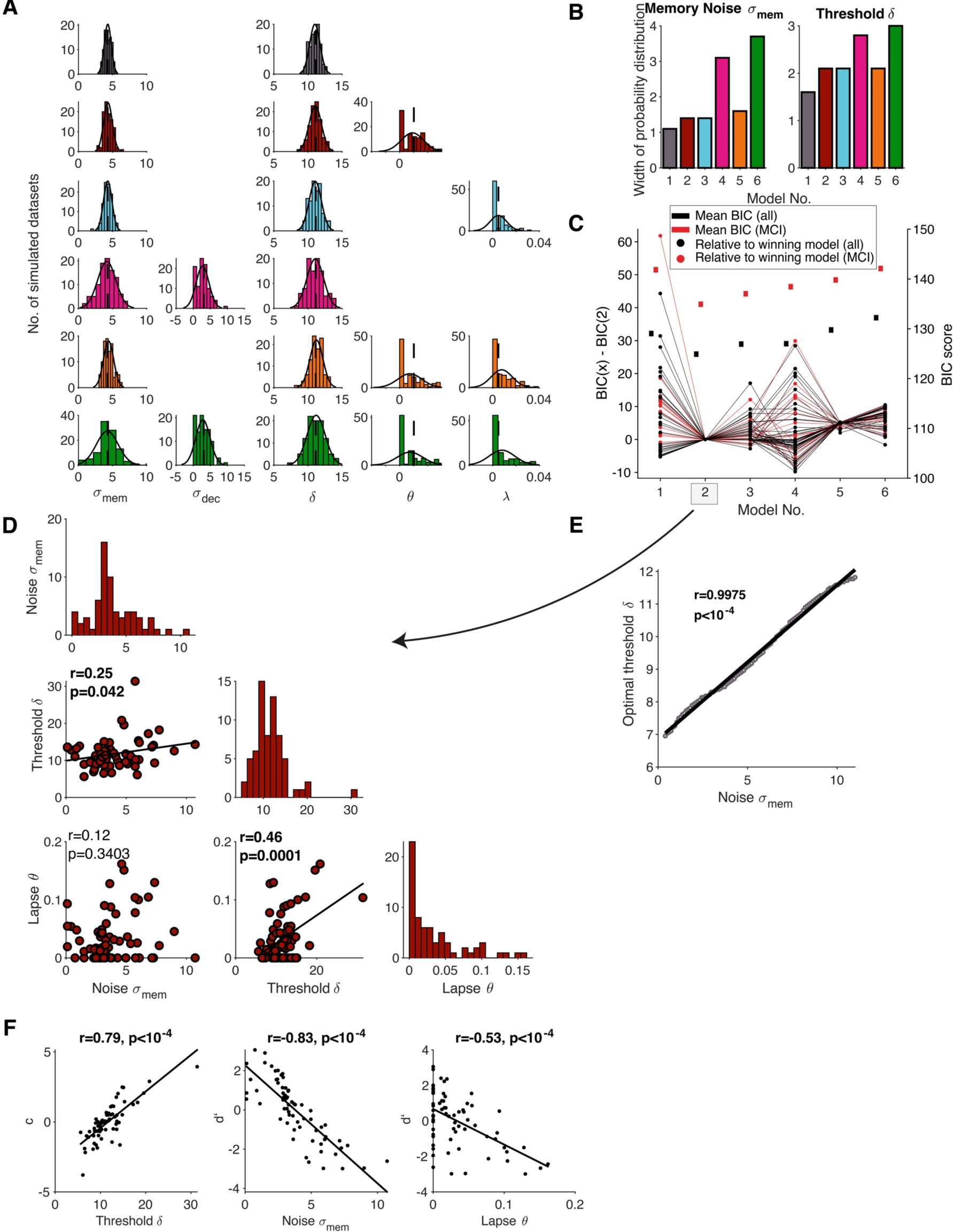
Model validation and comparison. **(A)** Histograms of fitted parameters for simulated data sets (models 1-6, colored), the corresponding fit of a normal density function (solid black line) and parameter levels at which the data sets were simulated (dashed black line). **(B)** Width of the probability density function for fitted model parameters that are included in all candidate models (memory noise and decision threshold) at half maximum. Models 4 and 6 (including decision noise as a free parameter) showed the strongest limitations in the recoverability of memory noise and threshold. **(C)** Comparison of the candidate models 1-6 from the mean BIC scores and the BIC score relative to the winning model for MCI patients (red) and the entire sample (black), indicating superiority of Model 2. **(D)** Intercorrelation and histograms of fitted model parameters for Model 2 across all subjects (N=67). **(E)** Memory noise levels and corresponding optimal threshold fits are strongly positively correlated in simulated data showing an optimality interdependence of the two parameters. **(F)** Correlations of fitted model parameters (N=67) with standard SDT measures of criterion (c) and sensitivity (d’) further serve as validation of fitted model parameters.

**Supplementary Figure 4.**
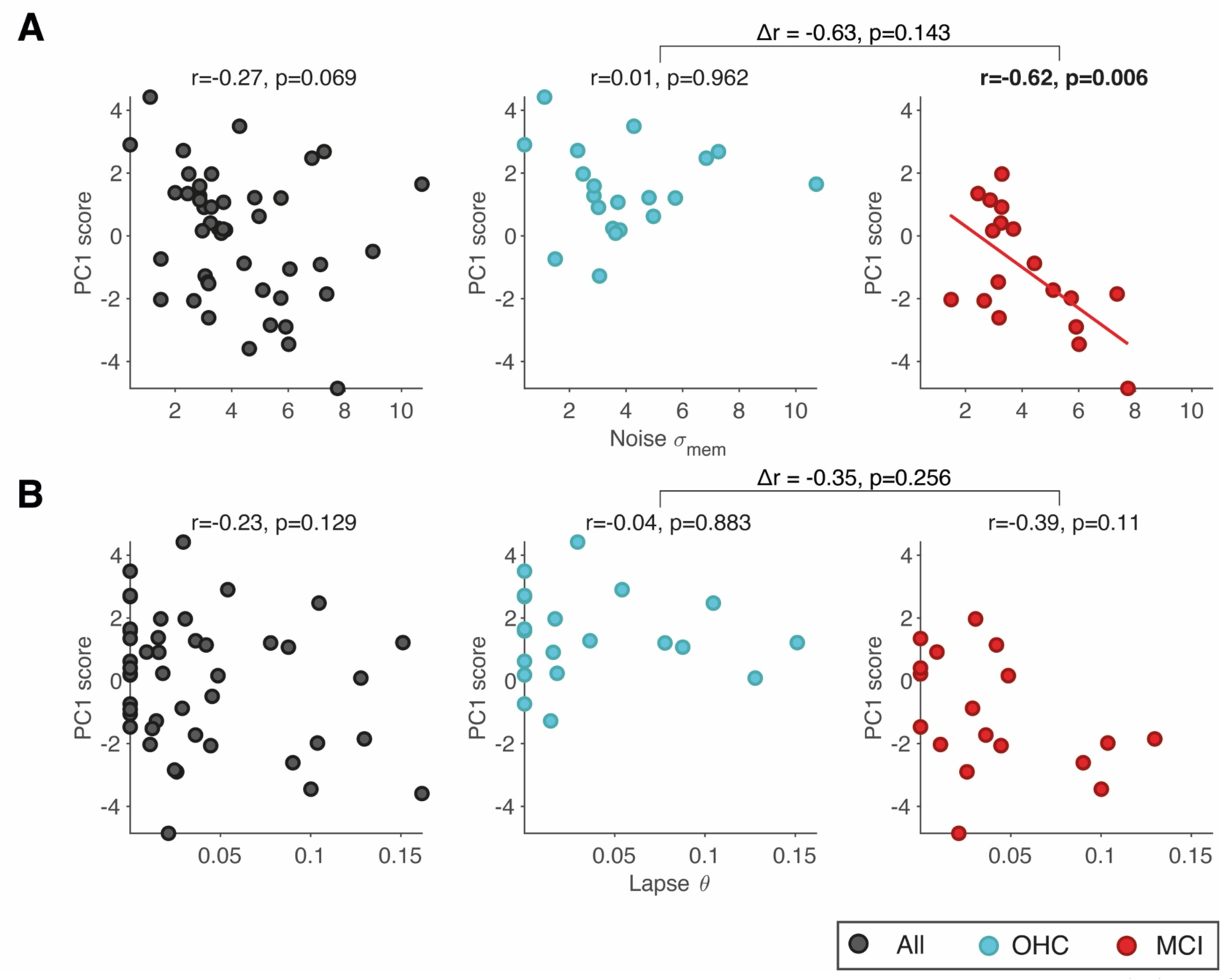
Correlation analyses with cognitive integrity (PC1) scores. Correlation of PC1 scores with **(A)** noise parameter fits and **(B)** lapse parameter fits. In all panels, circles represent individual subjects within the group of older participants. Correlations and correlational statistics are reported for all older subjects (N=45, black, *left*), OHC (N=20, blue, *middle*) and MCI (N=18, red, *right*) separately. Linear regression fit is shown for statistically significant correlations only. Differences in correlations between OHC and MCI are shown on top of the square brackets, p-values refer to two-sided permutation tests.

**Supplementary Figure 5.**
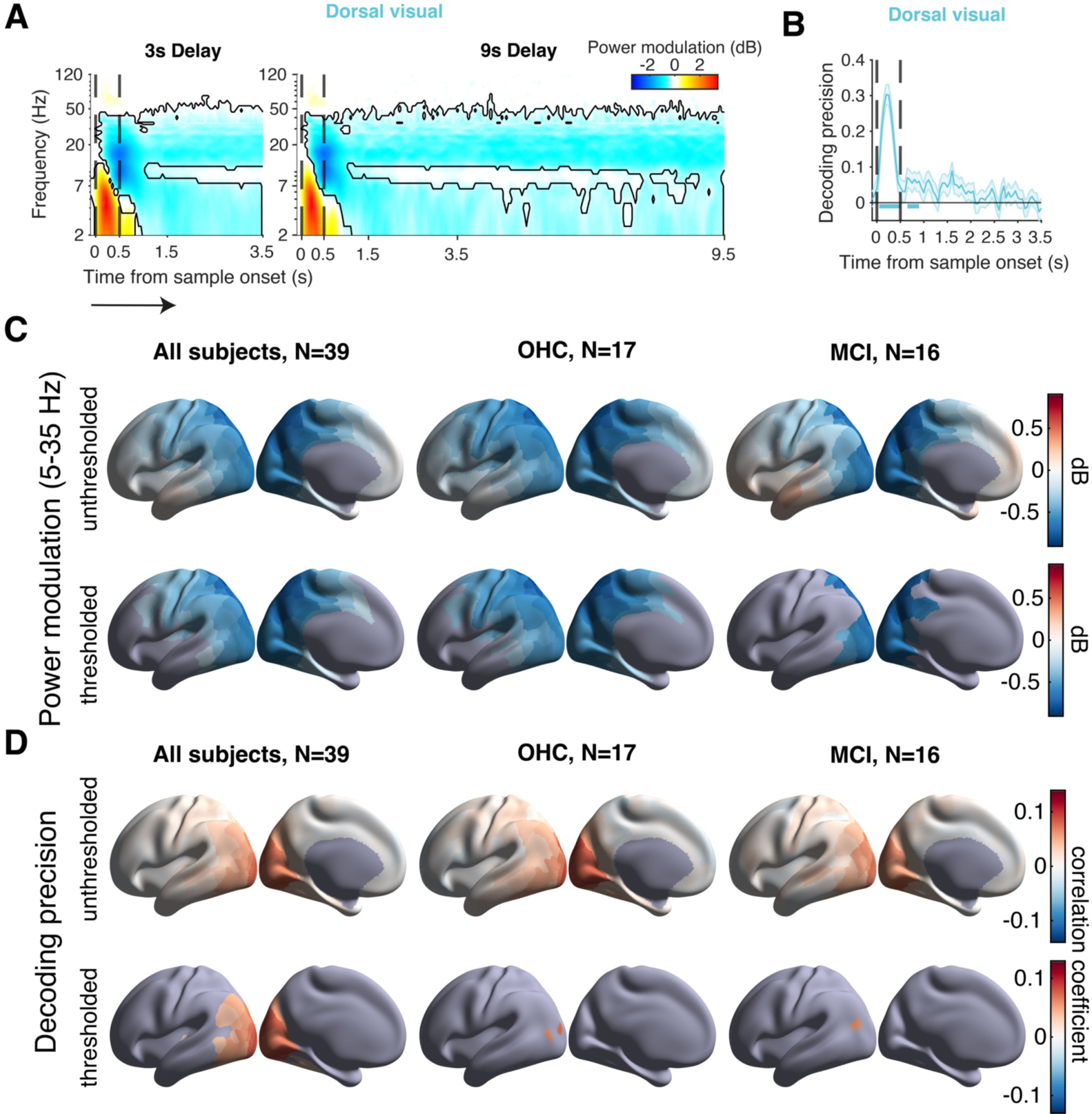
Power modulation & decoding precision on longer delay trials and brain maps for fine-grained cortical parcellation. **(A)** Task-induced power modulations for dorsal visual clustered brain region (see Table 2) for 3s delay (including first three seconds of trials with a 9s delay duration) and 9s delay show the persistence of power suppression with relative omission of alpha frequencies beyond the first second of delay (all subjects, N=39). Black contouring refers to significant within-subjects power modulation (cluster-based permutation test, *p*<0.05). **(B)** Time courses of vertex-based sample decoding precision in dorsal visual cortex (power values from 5-35 Hz; all older subjects, N=39; mean ± s.e.m.) during 3s of delay. Times of sample on- and offset are marked by vertical black dashed lines. Horizontal solid lines in respective color of brain cluster indicate significant decoding precision (cluster-based permutation test, *p*<0.05). Although dorsal visual showed robust encoding when considering only the first delay second of all trials, no significant decoding precision could be demonstrated throughout the 3s delay duration. **(C)** Fine-grained parcellated brain maps (N=180 ROIs) of unthresholded (*top*) and FDR-corrected (*bottom*) low-frequency (5-35 Hz) power modulation during first second into delay duration (significance threshold, *p*<0.05). **(D)** Fine-grained parcellated brain maps (N=180 parcels) of unthresholded (*top*) and FDR-corrected (*bottom*) decoding precision during first second into delay duration (significance threshold, *p*<0.05).

**Supplementary Figure 6.**
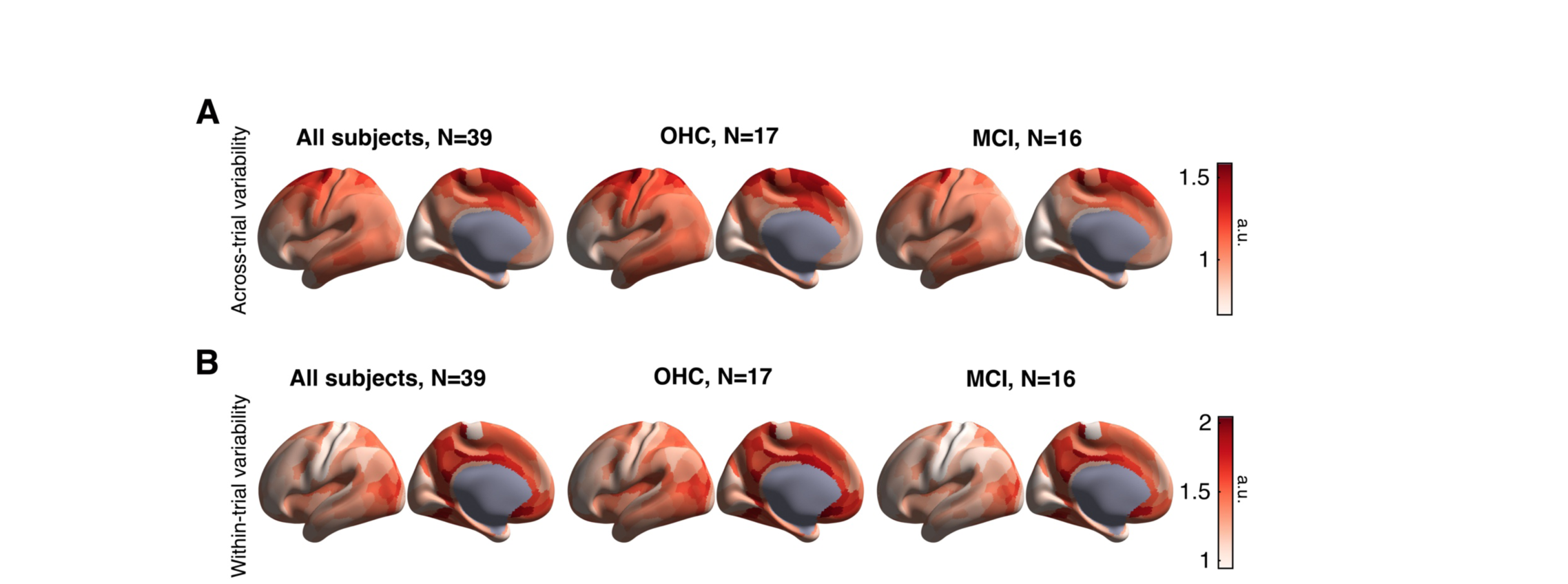
Cortical distributions of variability of task-related MEG power. **(A)** Maps of across-trial variability in power modulations during 1s of delay duration in the frequency range of 5-35 Hz. **(B)** As A, but for within-trial variability in power modulations.

## Notes

### Competing Interest Statement

The authors have declared no competing interest.

